# Pathogenesis and immune response to respiratory coronaviruses in their natural porcine host

**DOI:** 10.1101/2024.11.08.622602

**Authors:** Ehsan Sedaghat-Rostami, Brigid Veronica Carr, Liu Yang, Sarah Keep, Fabian Z X Lean, Isabella Atkinson, Albert Fones, Basudev Paudyal, James Kirk, Eleni Vatzia, Simon Gubbins, Erica Bickerton, Emily Briggs, Alejandro Núñez, Adam McNee, Katy Moffat, Graham Freimanis, Christine Rollier, Andrew Muir, Arianne C Richard, Nicos Angelopoulos, Wilhelm Gerner, Elma Tchilian

## Abstract

Porcine respiratory coronavirus (PRCV) is a naturally occurring pneumotropic coronavirus in the pig, providing a valuable large animal model to study acute respiratory disease. PRCV pathogenesis and the resulting immune response was investigated in pigs, the natural large animal host. We compared two strains, ISU-1 and 135, which induced differing levels of pathology in the respiratory tract to elucidate the mechanisms leading to mild or severe disease. The 135 strain induced greater pathology which was associated with higher viral load and stronger spike-specific antibody and T cell responses. In contrast, the ISU-1 strain triggered mild pathology with a more balanced immune response and greater abundance of T regulatory cells. A higher frequency of putative T follicular helper cells was observed in animals infected with strain 135 at 11 days post-infection. Single-cell RNA-sequencing of bronchoalveolar lavage revealed differential gene expression in B and T cells between animals infected with 135 and ISU-1 at 1 day post infection. These genes were associated with cell adhesion, migration, and immune regulation. Along with increased IL-6 and IL-12 production, these data suggest that heightened inflammatory responses to the 135 strain may contribute to pronounced pneumonia. Among BAL immune cell populations, B cells and plasma cells exhibited the most gene expression divergence between pigs infected with different PRCV strains, highlighting their potential role in maintaining immune homeostasis in the respiratory tract. These findings indicate the potential of the PRCV model for studying coronavirus induced respiratory disease and identifying mechanisms that determine infection outcomes.

**Author summary:** Understanding how our immune system reacts to respiratory viruses, like SARS-CoV-2, is crucial to developing better treatments. While most COVID-19 infections are mild, some cases lead to severe lung damage, but we do not fully understand why. To study this, we used pigs, which respond more like humans compared to small animals, to explore how the immune system deals with respiratory coronaviruses. We tested two porcine respiratory coronavirus strains that caused different levels of lung damage. The more severe strain triggered a strong immune response and high inflammation, leading to lung pathology similar to that seen in severe COVID-19 cases. By contrast, the milder strain caused a balanced immune response, including more regulatory T cells that help control inflammation. We also found changes in genes related to antibody-producing cells, which may be important for controlling respiratory pathology. Interestingly, changes in immune responses and gene expression lasted long after the virus was cleared, potentially making individuals more vulnerable to future infections - similar to the “long COVID” symptoms seen in people. We propose that this pig model could help us study coronavirus-induced lung damage and test new therapies to prevent severe disease.

## Introduction

Respiratory coronaviruses (CoV) are a significant threat to global health and have caused three major human epidemics of severe respiratory disease since 2003, including the recent SARS-CoV-2 pandemic. In each case, CoV emergence in humans has been associated with zoonotic transmissions from animals. Many livestock species are infected with CoVs causing variable morbidity and mortality, associated with lung disease and immunological impairment, leading to substantial economic losses [1, 2]. Porcine respiratory CoVs (PRCVs) are globally endemic in pigs. PRCV is a naturally occurring spike deletion variant of the *Alphacoronavirus* Transmissible Gastroenteritis Virus (TGEV) [3, 4]. TGEV formerly caused enteric disease with high mortality in neonatal piglets, but since the emergence of PRCV, the impact of TGEV has been greatly reduced due to cross protective immunity [5, 6]. PRCV infection is usually subclinical with most cases being mild unless secondary infections occur.

Previously it was suggested that PRCV infection with ISU-1 strain induces pulmonary pathology similar to that of SARS-CoV [7]. However, the lung pathology was mild unless potentiated with corticosteroids [7]. Given that pigs are large, long-lived natural hosts for CoV with respiratory anatomy and physiology similar to humans, we investigated the effects of different PRCV strains *in vivo.* We showed varying degrees of lung pathology: PRCV 135 strain caused a prominent pneumonia similar to human SARS-CoV-2, with viral tropism and damage to the bronchial and alveolar cells; in contrast, the pathological changes driven by ISU-1 infection were very mild [8]. Although PRCV strains 135 and ISU-1 replicated equally well in continuous porcine cell lines *in vitro,* only the more pathogenic 135 strain was able to replicate in *ex vivo* tracheal organ cultures, as well as *in vivo* in the trachea and lung [8]. PRCV uses aminopeptidase N (APN) as a receptor, although recent studies suggested another host entry or host restriction factor might be involved [9–11]. Differences in the Spike and gene 3 sequences between the strains maybe also responsible for the variation in replication patterns [12].

Mouse, ferret, hamster and non-human primate models have been developed to study SARS-CoV-2. However, these species are not natural hosts for the virus, and it remains uncertain which model most accurately mimics human disease [13–15]. In contrast, the pig is a natural host for several porcine CoVs including pneumotropic viruses [1, 16, 17]. As with SARS-CoV-2, PRCV infection can be mild or asymptomatic, but in some instances can lead to pneumonia. Better understanding of the mechanisms that lead to mild or severe disease is essential for developing new control strategies for emerging severe coronavirus diseases of livestock or humans. Here we used infection with the less and more pathogenic PRCV strains ISU-1 and 135, respectively, for in depth mechanistic evaluation of the pathogenesis, virology, and immune responses of this important family of viruses. We performed high resolution analysis of early and late systemic and mucosal immune responses using multi-parameter flow-cytometry to reveal how effector functions of the cellular and humoral responses mediate lung pathology. We also performed scRNA-seq on cells from bronchoalveolar lavage (BAL), as we previously did to characterise changes during influenza infection and respiratory immunisation [18]. We used the scRNA-seq data to interrogate early (day 1) and late (day 20) transcriptional changes following infection with ISU-1 and 135 PRCV strains. We propose that the pig is an appropriate and powerful model for understanding immunity to CoV in a natural large animal host, particularly the early innate and subsequent adaptive immune response at the site of infection, which are critical for protective immune responses and vaccine efficacy.

## Results

### Viral load and respiratory pathology following PRCV ISU-1 and 135 infection

Forty pigs were inoculated intranasally/intratracheally with 1 x 10^7 PFU of either PRCV ISU-1 or 135 strains. Five pigs from each group were humanely culled at 1, 5, 11, and 20 days post infection (DPI) **(Figure 1A)**. The quantity of infectious virus present in daily nasal swabs was determined by plaque assays. Peak viral shedding occurred between 2-6 DPI for both ISU-1- and 135-infected animals, followed by a rapid decline from 6 DPI. Viral shedding of 135- infected animals was significantly higher on 1, 4, 5, and 6 DPI compared to ISU-1-infected animals **(Figure 1B).** Total viral shedding over the time course of infection was also significantly greater in the 135-infected animals as determined by the area under the curve **(Figure 1C**).135-infected animals exhibited higher viral load in tracheal swabs, tracheal tissue, bronchoalveolar lavage (BAL) and lung, with a maximum level detected at 5 DPI and return to baseline at 11 DPI **(Figures 1D-G)**. Infectious virus was detected in the eyelid of the 135 group (4/5 pigs), which was comparable with studies reported from SARS-CoV-2 (**Figure 1H**) [19] A section of skin from the cheek (masseter region) was evaluated but no infectious virus was isolated (data not shown) indicating that the observed replication of 135 is specific for the conjunctiva rather than skin.

**Figure 1.**
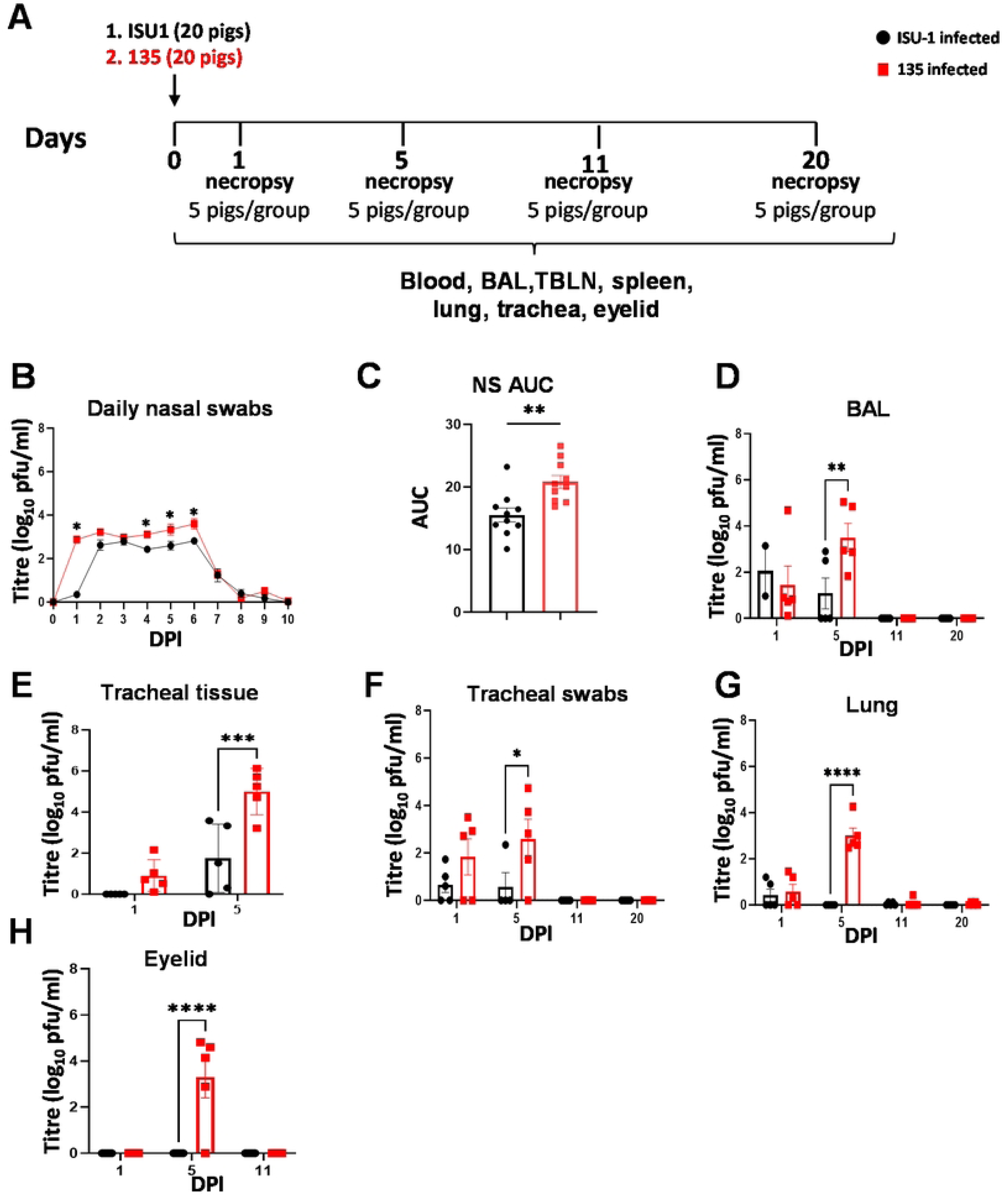
Experimental design and virus load in tissues after in vivo inoculation with PRCV strains ISU-1 and 135. (A) Forty pigs were infected with 1 x 10^7^ PFU of PRCV ISU-1 (black) or 135 (red) by the intratracheal/intranasal route (IN/IT). At 1, 5, 11 and 20 days post infection (DPI) five animals per group were culled. (B) Viral load in daily nasal swabs (NS). (C) Significant differences in nasal shedding determined by the area under the curve (AUC). (D) Bronchoalveolar lavage (BAL) fluid, (E) trachea tissue, (F) tracheal swabs, (G) accessory lung lobe and (H) eyelid were determined by plaque assay. Each point represents one animal with error bars defining SEM. Statistical differences at each time point were analyzed using unpaired t test with Welch’s correction or two-way ANOVA with a Bonferroni’s test for multiple comparisons. Asterisks represent significant changes (*p < 0.05, **p < 0.01, ***p < 0.001, ****p < 0.0001).

At necropsy, pulmonary consolidation was only observed in the 135 group, with an average of 1.2% lung areas affected at 1 DPI, 11.2% at 5 DPI, a reduction by 11 DPI to 5.8%, and absence of overt consolidation at 20 DPI (**Figure 2A**). Only the cranial and middle lung lobes were affected. No gross lung lesions were noted in the ISU-1 group throughout the study period. Using immunohistochemistry, viral antigen (nucleocapsid) was detected in the nasal turbinate (respiratory and olfactory aspects of nasal mucosa) of both groups at 1 DPI, with higher levels in 135-infected animals and consistently remained elevated until day 5 DPI. Viral antigen was not detected at 11 and 20 DPI **(Figures 2B and D)**. At 5 DPI the ISU-1 infected upper respiratory tract had mild rhinitis and tracheitis, whereas 135-infected animals exhibited overt pathology in both upper and lower respiratory tracts compared to ISU-1 **(Figures 2B and C)**.

**Figure 2.**
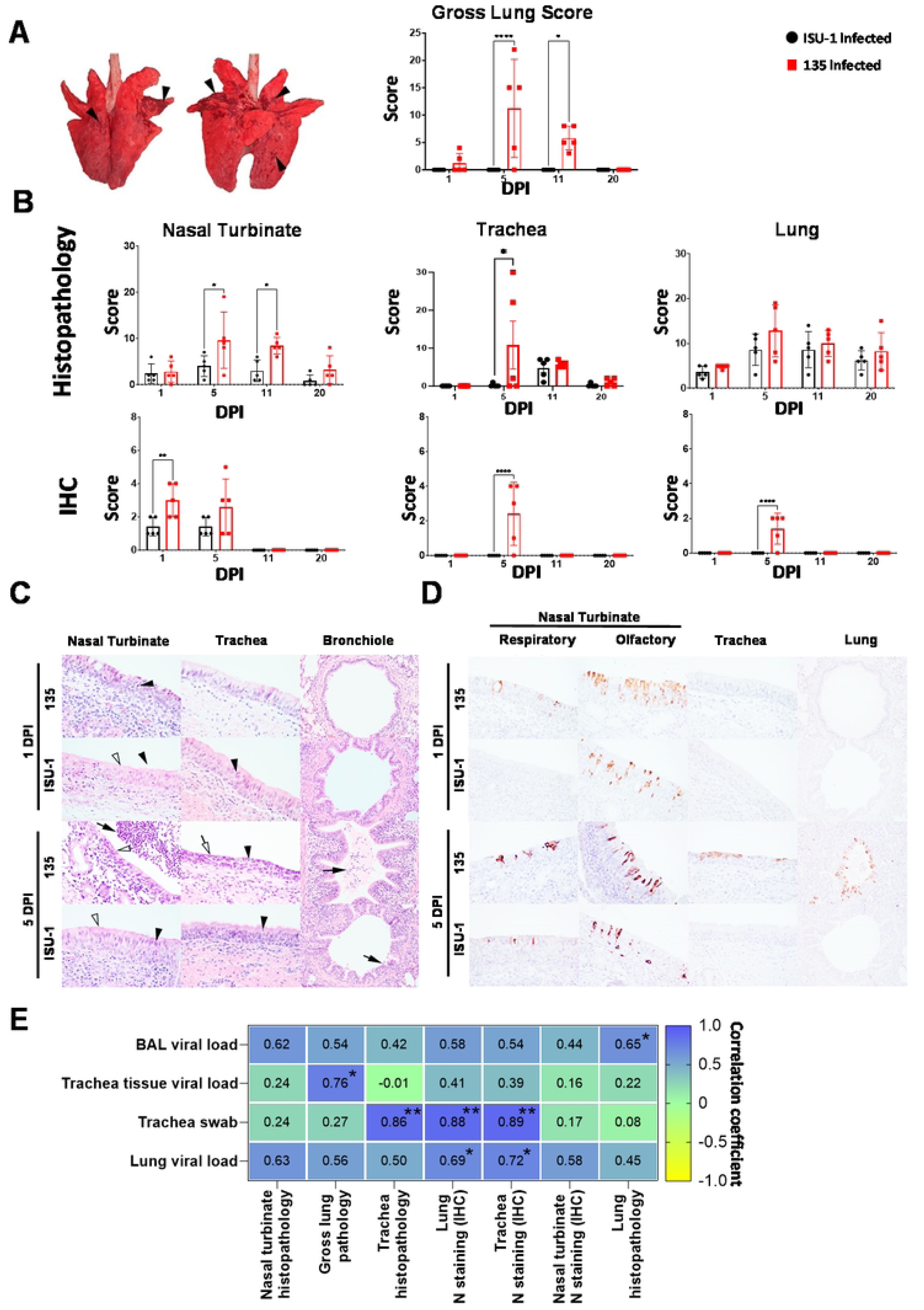
Development and progression of respiratory pathology following PRCV ISU-1 and 135 inoculations. **(A)** Representative gross lung pathology from 135 infected pig and percent area of lung consolidation for pigs infected with ISU-1 (black bars) or 135 (red bars) at 5 DPI. Arrowheads indicate areas of consolidation. Images are dorsal and ventral views **(B)** Histopathology and immunohistochemical scores for nasal turbinate, trachea, and lung for 1, 5, 11 and 20 DPI. **(C)** Representative histopathology from 1 and 5 DPI. **(D)** Representative viral immunohistochemistry from 1 and 5 DPI. Virus antigens were first detected in the nasal turbinate, with particular widespread labelling in the olfactory component, at 1 DPI in both groups, and was similar at 5 DPI. In the lower respiratory tract, virus antigens were only detected in the trachea and lung of 135 infected animals at 5 DPI, and these areas were colocalized with tracheitis and bronchopneumonia respectively. Images taken at 400x for nasal turbinate and trachea, 200x for lung. H&E and IHC were serial sections. Two-way ANOVA and Bonferroni’s multiple tests were performed. Asterisks represent significant changes (*p < 0.05, **p < 0.01, ***p < 0.001, ****p < 0.0001). **(E)** Spearman rank correlation between respiratory pathology and viral load at 5 DPI from ISU-1 and 135 infected animals. Asterisks represent significant changes (*p<0.05, **p<0.01).

At 5 DPI, animals infected with 135 developed bronchiolar exudation and epithelial damage, and within the alveoli there were frequent neutrophil accumulation (4/5 pigs) and oedema (2/5 pigs), all of which these lesions colocalised with virus antigen **(Figures 2B, C)**. In contrast, the ISU-1 group showed mild epithelial attenuation and rare neutrophilic exocytosis, and virus antigens were absent **(Figures 2B-D).** The overt respiratory pathology in 135 infected animals at 5 DPI was positively correlated with higher viral load in tracheal swabs, lung and BAL compared to the ISU-1 strain (**Figure 2E**).

At 11 DPI, virus antigens were absent in the nasal turbinate, trachea, and lung of both groups (**Figures 2B-D).** While virus antigens were not detected in the nasal turbinate in either group **(Figure 2B)**, mucosa epithelial damage with evidence of regeneration, persisted in the 135 group. In the trachea, both groups showed mild exudation, and the 135-group exhibited low levels of submucosal lymphoplasmacytic infiltrates. In the lung, both groups displayed low-level bronchiolar epithelial exudation and attenuation. Additionally, there was type II pneumocyte hyperplasia in 135 infected pigs (2/5 pigs).

At 20 DPI, virus antigens were not detected in the respiratory tissues. There were scattered lymphocytic infiltrates in the nasal turbinate submucosa for both 135 and ISU-1, and in the tracheal submucosal more commonly in 135 pigs (3/5) compared to ISU-1 (1/5). In the lung, both groups had BALT formation and rare bronchiolar exudation, and type II pneumocyte hyperplasia in a 135 infected pig (1/5 pigs). In other tissues including submandibular salivary gland, olfactory bulb, duodenum, or colon, no virus antigens or lesions were detected.

Pathology evaluation demonstrated that the PRCV 135 strain induced greater pathology in the respiratory tract than ISU-1 which coincided with peak viral load at 5 DPI, and with minimal to mild histopathological changes persisted until 20 DPI. Additionally, this was the first report of PRCV detection in eyelids of pigs, suggesting a potential virus dissemination from the respiratory tract to the eye.

### Antibody and B cell responses during ISU-1 and 135 infection

Antibody responses following ISU-1 or 135 inoculation were determined in serum, BAL and nasal swabs. Spike specific IgG and IgA were measured by endpoint titer ELISA and were detected in serum, BAL, and nasal swabs at 11 DPI and maintained at 20 DPI with significantly higher titers for IgG in the135-infected group compared to the ISU-1-infected group (**Figures 3A and B)**. IgA titers were lower compared to IgG. Serum from both 135- and ISU-1-infected animals cross-neutralized 135 and ISU-1 virus strains in addition to PRCV strains 137 and 310 and the parent TGEV strain **(Figure 3C)**. BAL titres were much lower compared to serum, likely due to the diluting effect of sampling. The histopathology in nasal turbinates and gross lung pathology at 11 DPI correlated with Spike-specific antibody responses in BAL, serum and nasal turbinates on the same day **(Figure 3D).**

**Figure 3.**
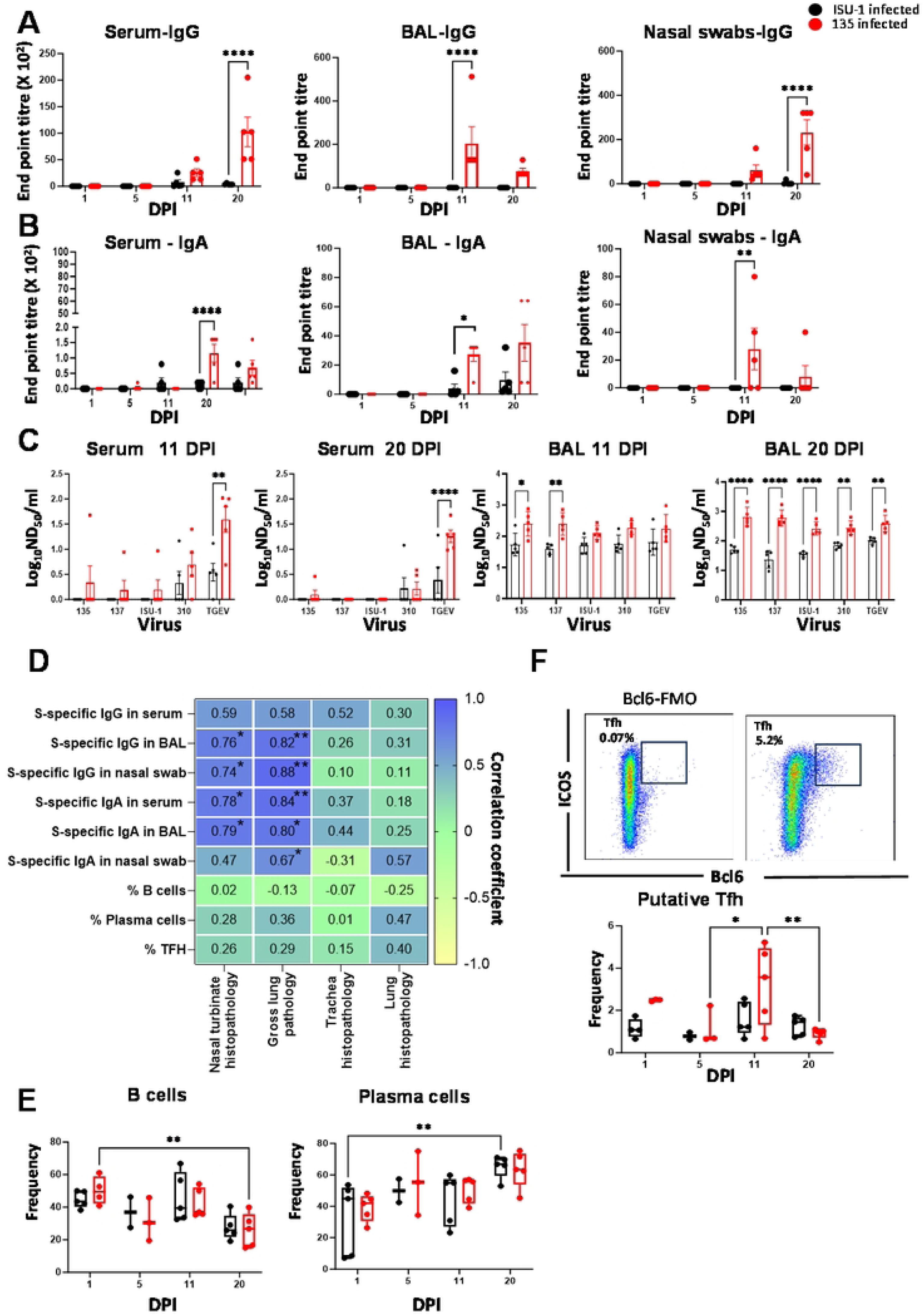
Antibody and B cell responses following ISU-1 and 135 infections. Spike specific IgG (A) and (B) IgA end-point titers were determined in serum, BAL and nasal swabs using ELISA. (C) Homologous and cross neutralizing antibody titers were determined using virus neutralization assay in Serum and BAL. Each point represents one animal with error bars defining SEM. Multiple Mann-Whitney test was performed changes between 2 groups. Asterisks represent significant (*p < 0.05, **p < 0.01, ***p < 0.001, ****p < 0.0001). **(D)** Spearman rank correlation between respiratory pathology and antibody responses from both ISU-1 and 135 infected samples at 11 DPI. Asterisks represent significant changes (*p<0.05, **p<0.01). **(E)** FACS staining was performed on BAL cells to identify total B and plasma cells. **(F)** Gating strategy for identifying BAL derived putative T follicular helper like cells and their proportions at 1,5, 11 and 20 DPI are shown. Each point represents single animal, and mid-line represents the mean. Two-way ANOVA and Bonferroni’s multiple tests was performed to compare temporal dynamics within each group and changes in frequency between 2 groups on single time point. Asterisks represent significant changes (*p<0.05, **p<0.01).

B cell responses in BAL were assessed by flow cytometry based on a previously established staining panel [20] **(Supl Fig S1A)**. Cells expressing CD79-α, PAX5, IRF4 and Blimp1 were defined as plasma blast/plasma cells. Within the IRF4 and Blimp1 double negative cells, the following subsets were further defined: proliferating B cells (Ki-67 positive), germinal centre B cells (Bcl6 and Ki-67 double positive) and naïve/memory B cells (Bcl-6 and Ki-67 double negative). There was a temporal shift from B cells towards plasma cells in the BAL, suggesting the differentiation of B cells to plasma blast/plasma cells in response to both infections **(Figure 3E)**. The shift coincided with a greater abundance of cells with a phenotype characteristic of T follicular helper cells (CD4^+^ICOS^+^Bcl-6^+^) in 135-infected pigs at 11 DPI compared to the ISU-1-infected group **(Figure 3F)**. Since the effector function of these cells has not been characterised in the porcine model, we refer to them as putative T follicular helper cells (Tfh). A respiratory resident CD4 T cell subset with phenotypic and transcriptional characteristics resembling both Tfh and tissue resident cells has been identified in a murine model and was shown to play a crucial role in inducing and maintaining memory B cells and CD8 T cells specific to influenza antigens [21].

In summary a stronger antibody response was detected in the serum, BAL and nasal swabs of 135 infected animals compared to ISU-1, which was dominated by IgG. Serum antibodies cross neutralized all PRCV strains and TGEV. For the first time we identified putative Tfh in porcine BAL indicating a role in regulating local humoral responses.

### Cytokine T cell responses in BAL, TBLN and spleen following ISU-1 and 135 infection

As T cells are crucial for control of virus replication, we examined CD4 and CD8β T cell responses during ISU-1 or 135 infection. We analysed cytokine producing cells in BAL, tracheobronchial lymph nodes (TBLN) and spleen by ELISpot and intracellular cytokine staining (ICS, **Supl Fig S1B**) following stimulation with 135 or ISU-1 live viruses, or overlapping pools of peptides covering the Spike (S) protein as previously described [8]. An increased number of Spike-specific IFN-γ-secreting cells was detected by ELISpot in BAL of 135-infected animals at 20 DPI, compared to ISU-1-infected animals **(Figure 4A, Supl Fig S2)**. Similarly greater proportions of 135- and S-specific CD4 IFN-γ-secreting cells were observed by ICS at 20 DPI, while 135- and S-specific TNF-secreting CD4^+^ T cells were significantly higher at 11 DPI **(Figure 4B)**. The 135-specific CD8 T cells exhibited the highest IFN- γ production at 11 DPI and were significantly more abundant in 135-infected animals compared to ISU-1-infected animals **(Figure 4C)**. The percentage of 135- specific IFN-γ- and TNF-secreting CD8^+^ T cells, as well as 135- and Spike-specific IFN-γ ELISpot responses, were associated with gross lung pathology and histopathology in nasal turbinates at 11 DPI (**Figure 4D**). No differences in proportions of CD8^+^ TNF- or CD4 and CD8 IL-2 producing cells were detected between groups (**Supl Fig S2A**).

**Figure 4.**
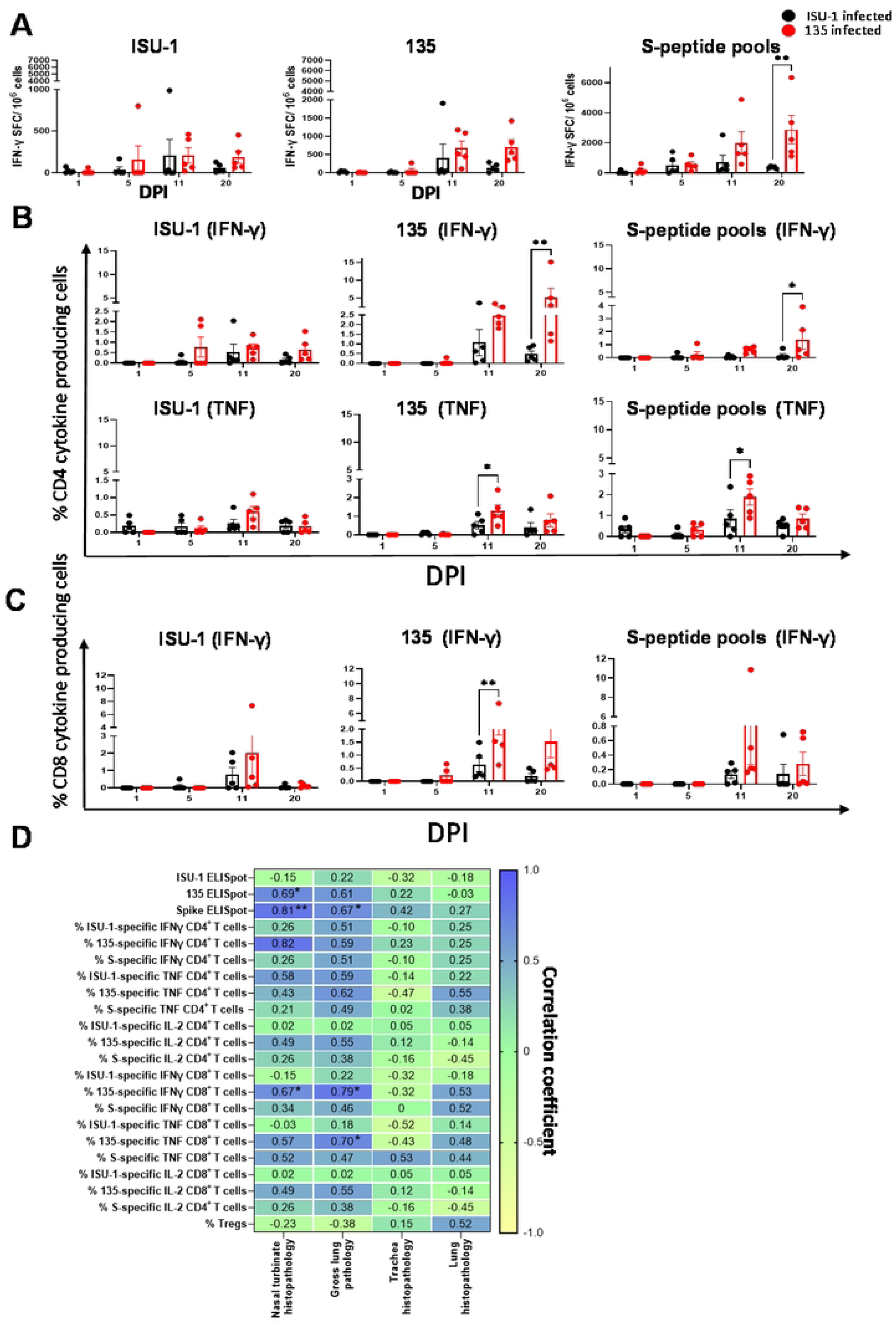
Cytokine T cell responses in BAL following ISU-1 or 135 infections. **(A)** IFN-γ secreting spot forming cells (SFC) were enumerated by ELISpot following *ex-vivo* stimulation with ISU-1, 135 or spike peptide pools. The frequency of IFN-γ and TNF secreting CD4 **(B)** and IFN-γ secreting CD8 **(C)** T cells were determined using intracellular staining following stimulation with ISU-1, 135 or peptide pools covering the Spike protein. Each point represents single animal. The line on each bar represent mean with SEM. Each point represents single animal and mid-line in the middle represent the mean. Two-way ANOVA and Bonferroni’s multiple tests were performed. Asterisks represent significant changes (*p < 0.05, **p < 0.01, ***p < 0.001, ****p < 0.0001). **(D)** Spearman rank correlation between respiratory pathology and T cell responses from both ISU-1 and 135 infected samples at 11 DPI. Asterisks represent significant changes (*p<0.05, **p<0.01). Asterisks represent significant changes (*p<0.05, **p<0.01 pathology from both ISU-1 and 135 infected 005).

Extending ELISpot and ICS to TBLN and spleen showed greater proportions of cytokine producing cells in 135-infected animals **(Supl Fig S3 and S4)**. IFN-γ ELISpot responses revealed significantly higher cytokine secretion in 135-infected animals at 11 DPI in TBLN and spleen **(Supl Fig S3A and S4A)**. ICS in TBLN demonstrated a higher proportion of virus- or Spike-specific IFN-γ and TNF-secreting T cells at 11 DPI or 20 DPI, and Spike-specific cells at 11 DPI in 135-infected animals (**Supl Fig S3B**). CD4 and CD8 IL-2-producing cells were also detected in TBLN with a significantly higher proportion in 135 infected animals at 20 DPI **(Supl Fig S3C**).

Taken together these data indicate that 135 infections induced greater IFN-γ, TNF and IL-2 T cell responses in BAL, TBLN and spleen at 11 DPI and 20 DPI compared to ISU-1 infection. The greatest response was in the BAL with high frequencies of CD4 IFN-γ- and TNF-producing cells.

### Single cell RNA-seq analyses from BAL cells

Since BAL cell preparations are predominantly composed of alveolar macrophages, we applied cell sorting to obtain matching proportions of macrophages, CD4 T cells, CD8 T cells, and B cells (12,000 cells each) from 4 animals per condition (ISU-1, 135) and 2 time points (1 DPI and 20 DPI), resulting in a total of 16 samples [18]. Cell partitioning and library preparation were performed separately for each sorted and re-pooled BAL sample.

To compare 135 and ISU-1 infection at each time point, samples were sequenced and analysed in DPI-specific batches. Graph-based clustering identified 19 and 20 distinct cell clusters in the 1 DPI and 20 DPI datasets, respectively. In the 1 DPI dataset, cluster 18 was of poor-quality and was excluded from further analysis. In the 20 DPI dataset, cluster 19 (containing a mixture of non-conventional T cells and macrophages) was excluded because almost all cells in this cluster originated from sample 12. In all other respects sample 12 showed patterns of cluster membership consistent with the other samples.

Some clusters were additionally split into subclusters to separate CD4 from CD8 T cells, non-conventional T cells from NK cells, or macrophages from monocytes, creating a final dataset with 39,679 and 149,069 cells for annotation in the 1 DPI and 20 DPI datasets, respectively. To label each cluster with conventional cell-type names, we used a combination of marker genes characteristic of each cluster based on a list of commonly defining marker genes. (**S1 Table**). From this information, each cluster was manually annotated with an established cell-type name for the 1 DPI and 20 DPI datasets (**Figure 5A, Supl Fig S5A**). In the 1 DPI dataset, subclusters 9b, 14b, and cluster 19 were enriched for NK cell-typical genes. The same applied to subcluster 2a and cluster 16 in the 20 DPI data set. Given that our cell sorting strategy did not include NK cells, these clusters were excluded to avoid a misrepresentation of these otherwise larger cell populations.

**Figure 5.**
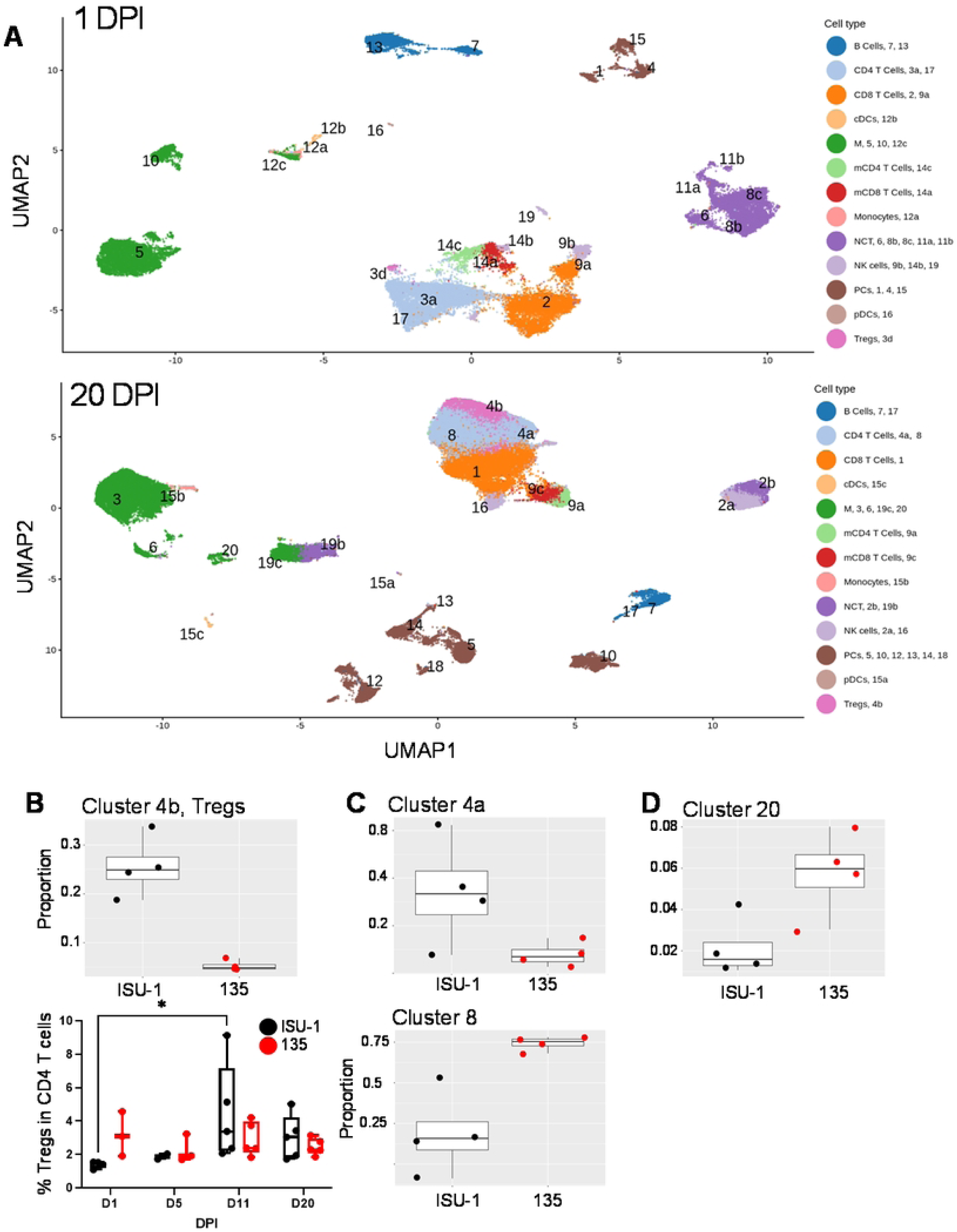
UMAP clustering and differential abundance analysis of BAL immune cells. **(A)** Uniform manifold approximation and projection (UMAP) visualization of all quality-controlled cells from pig BAL scRNA-seq analysis, colored by cell type and annotated with cluster numbers. Top: DPI 1 data set, Bottom: DPI 20 data set. Non-conventional T cells (NCT), Macrophages (M), Mitotic CD8 T Cells (mCD8 T Cells) **(B)** Proportions of Tregs (cluster 4b) at 20 DPI by scRNA-seq in CD4 T cells (top) and over the time course by flow cytometry (bottom). **(C)** Proportions of further cell clusters with significant differences between 135-infected and ISU-1-infected pigs within the CD4 T cell population at 20 DPI. **(D)** As in (C) but within macrophages.

### Differential abundance testing

We next tested the scRNA-seq data for differential abundance of cells within the sorted cell types (CD4 and CD8 T cells, B cells, macrophages) of ISU-1- versus 135-infected animals at each DPI (**S2, S3 Tables, Figures 5B-D**). No differences were detected at 1 DPI, but at 20 DPI pigs infected with the ISU-1 strain had significantly higher proportions of cells in cluster 4b, annotated as Tregs, within the sorted CD4^+^ T cell population (adjusted p=1.1×10^−4^, **Figure 5B** top). Tregs were further investigated by flow cytometry, based on a CD4^+^CD25^high^Foxp3^+^ phenotype (**Supl Fig S6A, Figure 5B** bottom). This analysis showed that in ISU-1-infected pigs, the proportion of Tregs was significantly higher at 11 DPI than at 1 DPI, but differences between 135 and ISU-1 infection were not statistically significant. Lack of precise concordance between scRNA-seq and flow cytometry Treg measurements likely reflects different definition parameters [18]. Nonetheless, our data indicate a greater proportion of CD4 T cells with a regulatory phenotype at later stages of ISU infection.

Differential abundance by scRNA-seq in Treg cluster 4b was mirrored by further differential abundances of the other clusters within the sorted CD4 T cell population **(Supl Fig S6B)**. Cluster 4a was also less abundant in 135-infected pigs (**Figure 5C**, top), while cluster 8 was more abundant (**Figure 5C**, bottom, **Supl Fig S6B**). While we cannot deduce whether these differences between PRCV strains result from expansion or reduction of the investigated clusters, it is of interest to look at key transcripts from clusters 4a and 8 to understand the subpopulations they represent. Cells in cluster 8 expressed higher levels of FCG3A, encoding CD16, but also IFITM1 or CD40LG, suggesting that cells in this cluster are capable of a variety of immune functions. Cells in cluster 8 expressed higher levels of RPS genes (e.g. RPS7, RPS8, RPS15A, RPS20 and RPS27A), indicating high protein synthesis activity. Other highly expressed transcripts of cells in cluster 8, such as TMSB4X, TMSB10, RAC2, MYL6, S100A4 and CORO1A, are involved in cell migration, suggesting recent influx of these cells into BAL. In contrast, cluster 4a was characterised by transcripts nvolved in T cell activation, including ICOS, CD28, RIPO2R, FYN, FYB1, CD247 and IL18R1. Together, these results indicate that at 20 DPI, the BAL CD4 T cell population in 135-infected animals was dominated by a translationally active, migratory population, while that in ISU-1-infected animals had a greater abundance of regulatory and active helper T cell populations.

Further differences in abundance at 20 DPI were found in macrophage cluster 20 which were increased in 135-infected animals (**Figure 5D, Supl Fig S6B**). Prominent transcripts in this cluster revealed a mixed profile. For example, DOCK4, SDC2, CCDC88A, and CDC42BPA are involved in cytoskeletal reorganization and migration, suggesting either recent influx into the lung or increased cellular trafficking. Other highly expressed genes including ENPP1, PPARG, AGPS, and GPCPD1are linked to purinergic signalling and lipid metabolism. Together, this profile may point towards a role in resolving inflammation. This interpretation aligns with the development and progressive recovery from viral pneumonia in 135-infected pigs by 20 DPI (**Figure 2A, 2B**).

Overall, these data indicate that PRCV infection with strains of differing virulence changes the balance of macrophage and CD4 T cell subsets at the late stages of infection 20 DPI. The higher proportion of Tregs in ISU-1-infected pigs suggest a role in modulating the immune response, potentially contributing to milder lung pathology. In contrast, 135-infected animals showed greater abundances of T cells with high protein synthesis and involvement in cell migration, including inflammatory exudation in the bronchiole and alveoli. 135-infected pigs also had more macrophages with a potential role in tissue healing at 20 DPI.

### Differential expression (DE) analysis

To investigate differential gene expression between the 135 and ISU-1 infections at 1 and 20 DPI, we used edgeR to conduct a pseudo-bulk DE analysis within each cluster. The number of DE genes per cluster are summarised in **Supl Fig S7**.

To determine whether differentially expressed (DE) genes were unique or shared between cell types (macrophages, CD4 T cells, CD8 T cells, B cells), significantly DE genes with a false discovery rate below 0.05 and absolute log fold change higher than 0.6 were visualized using Venn diagrams for 1 and 20 DPI. (**Figure 6A and Figure 7A**, respectively). At 1 DPI, only a small number of genes were differentially expressed between 135 and ISU-1 infected pigs (**Figure 6A-B, S4 and S5 Tables**). Nearly all of them were lower in 135-infected pigs and most of them were detected in B cells or plasma cells.

**Figure 6.**
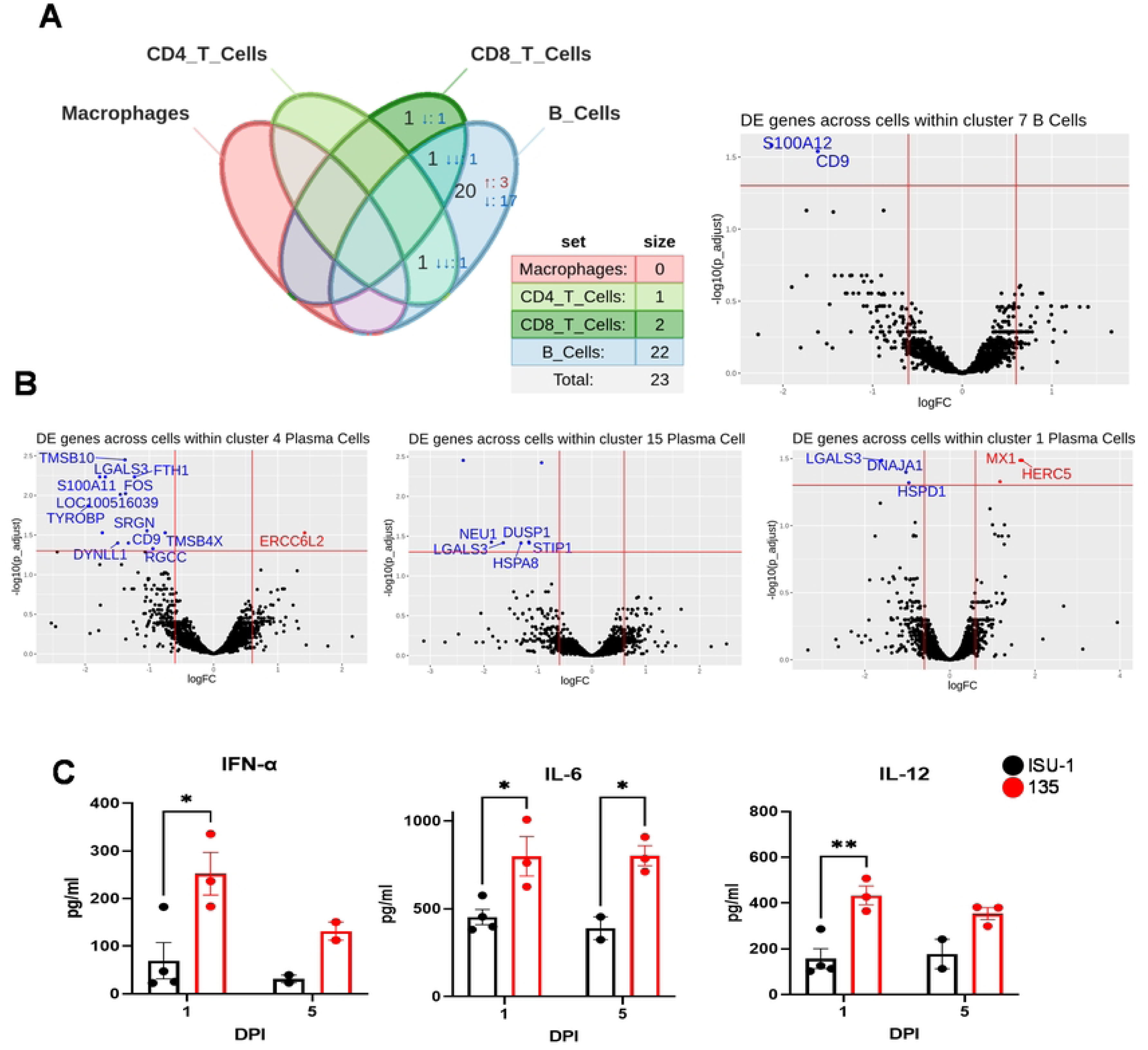
Differential gene expression and cytokine production comparing 135 and ISU-1 infected pigs 1DPI. **(A)** Venn diagrams shows the overlapping and unique differentially expressed genes for macrophages, CD4 T cells, CD8 T cells, and B cells at 1 DPI. Plasma cells were included in the B cells. Numbers represent differentially expressed genes, while the number of arrows indicates the number of cell types with differential expression. Red up arrows denote genes with significantly higher expression, and blue down arrows denote genes with significantly lower expression in 135-infected pigs. All genes shown have an adjusted p value (Benjamini-Hochberg) of less than 0.05 when comparing 135 with ISU-1. **(B)** Volcano plots of DE analysis results between 135- and ISU-1-infected pigs within cluster 1, 4, 15, 7 at 1 DPI, respectively. **(C)** BAL cells from ISU-1 (black) and 135 (red) infected pigs were stimulated overnight with the respective homologous virus strain. The supernatant was collected and Luminex assay was performed to measure the concentration of IFN-α, IL-6, IL- 12,

**Figure 7:**
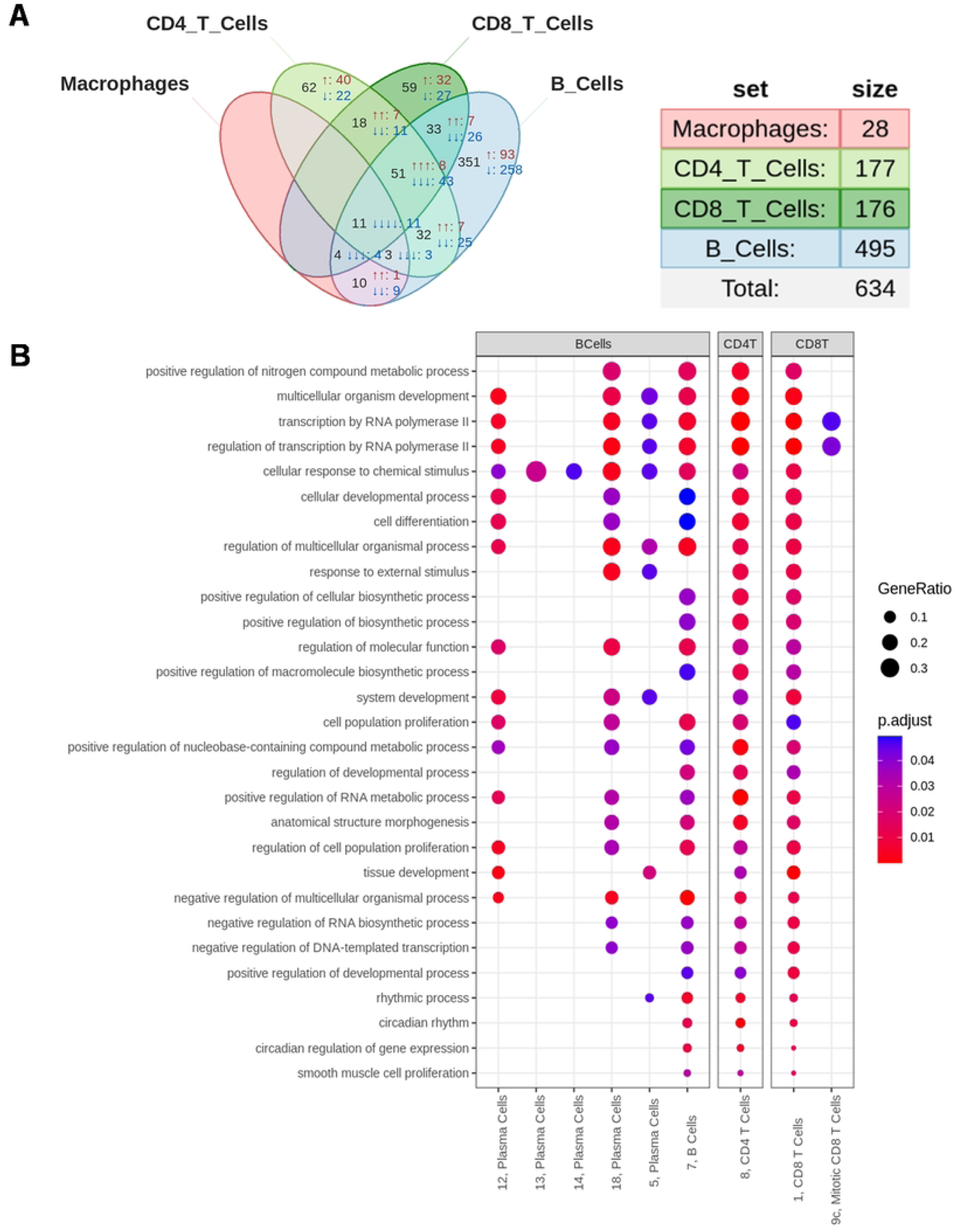
Differential gene expression at 20 DPI. **(A)** Venn diagram shows the number of overlapping and unique differentially expressed genes for macrophages, CD4 T cells, CD8 T cells, and B cells at 20 DPI. Plasma cells were included in the B cells. **(B)** Bubble plots presenting all enriched GO term shared across B cells, CD4 T cells and CD8 T cells populations at 20 DPI, based on all genes with an adjusted p value of less than 0.05. Bubble size represents the ratio of the number of DE genes associated with a specific GO term to the total number of DE genes identified in that cluster. Bubble colour represents the adjusted p-value. Terms are ranked according to gene ratio values within the 3 cell populations. Macrophage populations did not show significantly enriched GO terms. Clusters with no shared enriched GO terms are not shown.

Two genes were expressed at a significantly lower level in 135-infected animals in two cell types: NEU1 in CD4 T cells and B cells; and LGALS3 in CD8 T cells and B cells. NEU1 has the capacity to influence cell adhesion, which might alter cell-cell interactions of CD4 T cells and B cells in the BAL of 135-infected pigs [22]. Galectin 3, encoded by LGALS3, has been shown to negatively regulate T-cell activation [23]; lower expression of LGALS3 in CD8 T cells and B cells in 135-infected pigs may reflect increased activation at 1 DPI[23]. In addition to these shared DE genes, there were several DE genes unique for B cells, the majority of which were lower in the 135-infected pigs. Of note, most of these genes are derived from plasma cell clusters 4 and 15 (**Figure 6B**). Similar to NEU1, some of them are involved in cell adhesion and migration: TMSB10 and TMSB4X regulate actin polymerisation, while the tetraspanin CD9 is involved in the organisation of cell membranes. Further differentially expressed genes that were lower in 135-infected pigs included FOS and TYROBP, both of which are involved in immune cell signal transduction, while DUSP1 can dephosphorylate MAP kinases [24–26]. Of note, porcine BAL preparations contain high proportions of plasma cells within total B cells (**Figure 3D)** [20]. Their contribution to antibody production is currently unclear and some of these cells could also have regulatory functions [27].

Given the lower expression of genes related to cell adhesion and cell signalling, this could indicate reduced regulatory capacity of plasma cells in 135-infected pigs. Such postulated lower regulatory function may be linked to another experimental finding. We analysed early cytokine responses in BAL cell preparations using Luminex assays to measure IFN-α, IL-6, IL-12, IL-4, IL-1β, IL-8, TNF, and IL-10 at 1 and 5 DPI after *ex vivo* stimulation with virus homologous to the infection strain. BAL cells from 135-infected animals produced significantly higher concentrations of IFN-α, IL-6, and IL-12 in comparison with BAL cells from ISU-1-infected animals at 1 DPI **(Figure 6C and Supl Fig S9),** indicating stronger inflammatory responses in 135-infected pigs. This also aligns with higher expression of the antiviral MX1 gene in Tregs (cluster 3d, **Supl Fig S8**) and the plasma cell cluster 1 (**Figure 6B**).

In contrast to 1 DPI, at 20 DPI the number of genes differentially expressed between ISU-1 and 135 infected pigs was much higher (634 in total compared to 23 at 1 DPI), and comprised all cell types, with several DE genes shared between cell types (**Figure 7A, S6 Table**) and cell clusters (**Supl Fig S10**). Overall, these DE genes showed a very heterogeneous picture. For example, among the genes with lower expression in CD4 and CD8 T cells from 135-infected pigs were CD69 and CD28, involved in T cell activation and co-stimulation. Transcripts of the anti-apoptotic BIRC3 gene were also lower, suggesting a reduced capacity for stimulation or survival, but transcripts of the pro-apoptotic FAS gene were also lower. The dichotomy of simultaneously lower BIRC3 and FAS expression was also found in B cells and macrophages. Similarly, genes like ERBB4 and LRRK2, which are involved in cell activation [28], were both more highly expressed in CD4 and CD8 T cells of 135-infected pigs and contrasted the lower expression of genes encoding CD69 and CD28. The anti-apoptotic BCL2 gene was also expressed to a greater level in CD4 T cells of 135-infected pigs. This suggests that the two PRCV strains drive separate immune activation pathways, which are, however, often similar in their overall function.

We also performed Gene Ontology (GO) enrichment to investigate broader biological functions of the DE genes at 20 DPI (**S7, S8 Tables**). Several GO terms were shared between B cell clusters (including plasma cells), CD4 and CD8 T cells (**Figure 7B, Supl Fig S11**). Nearly all of those terms represented high-level biological functions like ‘multicellular organism development’, ‘transcription by RNA polymerase II’, ‘cellular response to chemical stimulus’, ‘regulation of molecular function’ or ‘cell differentiation’. This indicates that even 20 days after infection, when PRCV has already been cleared for 10 days (**Figure 1**) the more pathogenic 135 strain continues to impact gene regulation and cell metabolism in immune cells in the BAL. Of note, as on 1 DPI, B cells and the various plasma cell clusters showed the highest number of DE genes compared to other cell types (**Figure 7A, Supl Fig S7B**), indicating a hitherto unrecognised impact of highly pathogenic virus infection on gene regulation in local B cells and plasma cells.

In summary, these data suggest that the strong inflammatory processes triggered as early as 1 DPI have lasting consequences on the transcriptional landscape of immune cells in BAL, even after the virus has been cleared. Our data also indicate that differences in gene expression between pigs infected with the ISU-1 or 135 PRCV strains are most prominent in B cells during the early stage of infection. The absence of differentially expressed genes in macrophages suggests that this cell type responded similarly to both infections. Additionally, higher IL-6 and IL-12 cytokines in 135-infected animals may be associated with greater pulmonary lesions in these animals.

## Discussion

The spectrum of outcomes following SARS-CoV-2 exposure in humans is wide and considered to be largely determined by the host response. Most infections are asymptomatic or mild, indicating that effective host responses can successfully dampen viral replication and symptoms. However, the factors that precipitate severe disease are not fully understood. There is therefore a need for models that closely resemble human SARS-CoV-2. Currently mouse, ferret, hamster and non-human primate models have been developed to study SARS-CoV-2 but these are not natural hosts. In contrast pigs are natural hosts for PRCV, but little is known about the mechanism of pathogenicity and protective immunity in this species. Here we used infection with the weakly and strongly pathogenic PRCV strains ISU-1 and 135, respectively, for in-depth mechanistic evaluation of the pathogenesis and immune control of respiratory CoV.

Pathological examination revealed prominent respiratory lesions in 135-infected animals at 5 and 11 DPI which was still present at 20 DPI. This was characterised by a marked upper and lower respiratory pathology and with an associated high viral shedding in nasal swabs and viral load in trachea, lung and BAL at 5 DPI. Further, infectious virus was detected in the eyelids of a large proportion of 135 infected animals at 5 DPI, which suggest a similar pathogenesis as that reported in humans with SARS-CoV-2 infection [19, 29, 30]. While the virus tropism was not examined by microscopy, the ability of 135 to replicate in eyelid tissue, likely conjunctiva epithelium, may represent an understudied site of viral replication and potential route of both infection and transmission for respiratory CoV.

As well as being more virulent and pathogenic, 135 infection induced greater Spike specific IgG and IgA titers in serum, BAL and nasal swabs compared to ISU-1 infection. The increased antibody response was associated with a corresponding increase in plasmablast/plasma cells in the BAL. Additionally, we identified for the first-time cells with a Tfh phenotype which likely play a crucial role in maintaining local B cell and CD8 T cell memory [21].

Infection with 135 induced a higher proportion of virus specific IFN-γ and TNF secreting CD4 and CD8 T cells. The frequencies of cytokine secreting CD4 and CD8 T cells were greater in the BAL compared to TBLN and spleen in both groups. The highest frequencies of IFN-γ and TNF-secreting cells and the highest antibody titers were detected at 11 and 20 DPI, while the peak of lung pathology and viral load was at 5 DPI, implying that, along with the greater ability of 135 virus to infect more cells, the infection also resulted in a dysregulated and/or more vigorous innate immune response. Consequently, the presence of more virus antigen and this robust innate response may further drive a more vigorous adaptive immune response which may additionally contribute to the lung damage. This could be consistent with findings in SARS-CoV-2 patients where higher viral load was associated with increased disease severity along with robust activated CD4 T cells, activated CD8 TEMRA, hyperactivated or exhausted CD8 T cells and plasmablasts [31–33].

We therefore examined the early (1 DPI) and late (20 DPI) events in the BAL of animals infected with 135 and ISU-1 to identify gene expression changes that may lead to more severe or less severe inflammatory and pulmonary pathology. We analysed BAL as this is an easily accessible site for sampling in both humans and pigs and we have previously established the feasibility of performing single-cell transcriptomics with these samples in pigs [18]. Single-cell RNA-seq analysis of BAL cells revealed that the more pathogenic 135 strain shifts the balance of CD4 T cells away from Tregs (cluster 4b) toward translationally active CD4 T cells (cluster 8) at 20 DPI. The higher proportion of biosynthetic CD4 T cells in the 135-infected animals may have contributed to the increased antibody production, while the more abundant Tregs in ISU-1-infected pigs were associated with lower pulmonary lesions.

Differential expression analysis also indicated variation in gene expression between 135- and ISU-1-infected pigs at 1 DPI among genes related to cell migration and immune system processes. In 135-infected animals, higher expression of the antiviral gene MX1 in plasma cells, along with increased cytokine production of IL-6 and IL-12 in PRCV-stimulated BAL cultures, indicate a heightened inflammatory response, which might contribute to more severe pulmonary pathology. The higher concentration of IFN-α in 135-infected pigs may be attributed to higher viral load but is unlikely to be associated with the increased pulmonary pathology [34]. However, elevated IL-6 production has been associated with severe disease in COVID-19 patients, while IL-12 promotes Th1 differentiation and NK cell activation [35–37].

Most of the DE genes at 1 DPI were found in B cells, primarily in plasma cells. While the total number of DE genes was much higher at 20 DPI, with a high number in B and plasma cells, many were also shared between B cells and CD4 and CD8 T cells. A notable finding was that most of those DE genes were lower in cells from 135-infected pigs compared to ISU- 1-infected pigs. This may indicate a hitherto unnoticed role of B cells and plasma cells in local immune homeostasis in the lung. Regulatory functions of B cells and plasma cells have been described in a wide range of immune responses in mice and humans [27, 38]. Since B cells with regulatory functions do not form a defined cell lineage, no universal phenotype or master transcription factor is available for their identification. However, CD9, which was among the genes with lower expression in B cells and plasma cells of 135-infected pigs, has been described as a molecule associated with IL-10 competent regulatory B cells in mice [39].

GO analyses indicated that at 20 DPI genes DE between 135- and ISU-1-infected pigs are associated with high level cellular functions, such as gene transcription and cell matabolism. This applied to B cells (including plasma cells) but also CD4 and CD8 T cells. This suggests, that despite clearance of virus and a reduction in pathology at this time point, longer lasting effects caused by the two different PRCV strains are maintained. This may have significant implications for immune responses to subsequent viral or bacterial infections in the respiratory tract, commonly seen with “porcine respiratory disease complex” [40, 41]. This phenomenon may arise not only from the pathogens involved, but also by the underlying priming of the immune system by previous infections, despite successful clearance. This is reminiscent of post viral syndrome reported in humans, such as the long-term damage in COVID-19 patients, commonly known as “long COVID” [42]. The pig model may offer an opportunity to experimentally explore these effects by following PRCV infection with another infection, providing a platform to study post-viral syndrome changes.

Overall, the data suggest that the more pathogenic 135 strain leads to increased virus replication and triggers stronger, potentially damaging innate and adaptive immune responses, characterized by heightened inflammation, leading to more prominent respiratory pathology. In contrast, the less pathogenic ISU-1 strain induces a more regulated immune response, with more Tregs and controlled cytokine production, resulting in milder pathology. These results agree with similar studies in humans highlighting the importance of timely and appropriately scaled immune responses to SARS-CoV-2 and that early intervention with therapies that dampen immune responses, such as steroids, has proved crucial in preventing progression to severe disease [43–46].

Our data provide a reference map for changes in gene expression early in PRCV infection and identify potential targets for anti-inflammatory drugs. Candidate interventions could be tested in the pig PRCV model, which more accurately reflects pathological inflammation in the respiratory tract of humans. Our model will also contribute to determining how to achieve robust virus-specific immunity to clear virus infection without causing exacerbated inflammation and immunopathology, which is crucial for limiting life-threatening disease.

## Materials and Methods

### Viruses

The previously characterized PRCV strains 86/135308 (referred to as 135, GenBank accession number OM830318), ISU-1 (GenBank accession number OM830321), 86/137004 (referred to as 137, GenBank accession number OM830320), AR310 (referred to as 310, GenBank accession number OM830319) and TGEV strain FS772/70 were used (1). All viruses were grown and quantified in Swine Testis (ST) cells and titres expressed as the number of plaque-forming units per ml (PFU/ml).

## Ethics Statement

Animal experiments were approved by the ethical review processes of The Pirbright Institute and the Animal and Plant Health Agency (APHA) according to the UK Animals (Scientific Procedures) Act 1986 under project license PP7764821.

### Animal study

Forty outbred female pigs, aged 4-6 weeks and genetically composed of 1/4 Large White, 1/4 Landrace, and 1/2 Hampshire breeds, were included in the study. Prior to the challenge, all animals were assayed for the presence of Spike antibodies to ensure their PRCV free status and randomly assigned into two groups of 20 pigs. One week after acclimatization, one group were inoculated with 1 x 10^7 PFU PRCV 135 strain and the other with 1 x 10^7 PFU PRCV ISU-1 strain in a total of 4ml (2ml per nostril) using a mucosal atomisation device (MAD). Clinical parameters including demeanour, appetite, respiratory signs, sneezing, coughing, nasal and eye discharge, faeces consistency and rectal temperature were assessed. One ISU-1- and five 135-inoculated animals showed mild clinical signs between 2 and 5 DPI, which resolved rapidly and did not exceed a score of 2 out of maximum 19. Animals were humanely culled at 1, 5, 11, and 20 DPI with 5 animals sampled from each group at each time point for assessment of viral load, pathology and immune responses. Daily nasal swabs were collected for evaluation of viral shedding. At postmortem bronchoalveolar lavage (BAL), tracheobronchial lymph nodes (TBLN), and spleen were collected, mononuclear cells isolated and cryopreserved as previously described [47]. Because of a technical failure no spleen cells were available to perform ICS at 20 DPI. Viral load was also assessed in BAL, lung, trachea, tracheal swabs and eyelids.

### Lung gross pathology, histopathology, and immunohistochemistry

The percentage of pulmonary consolidation, including both dorsal and ventral aspects were assessed blindly by a veterinary pathologist at necropsy [47]. Formalin fixed tissues were processed by a routine histology method. Haematoxylin and eosin staining and immunohistochemistry (IHC) against PRCV nucleoprotein (N) using a monoclonal antibody were performed on serially sectioned formalin-fixed paraffin embedded tissues [47]. Positive control slides containing either TGEV or PRCV infected or uninfected cells, as well as pig tissues derived from PRCV-challenged experiments, were included in IHC experiments to confirm specificity of immunolabelling.

### Virological assays

The evaluation of infectious viral progeny involved the analysis of harvested BAL fluid, tracheal and nasal swabs obtained post-mortem, and daily nasal swabs. All samples, stored in Viral Transport Medium (VTM), were subjected to plaque assay in ST cells using a previously established protocol [47]. Tissues collected at post-mortem, including trachea, lung, cheek, and eyelid, were homogenized in phosphate buffer saline “a” (PBSa) using a Tissuelyser II (Qiagen) with a 5 mm stainless steel ball (Qiagen). The resulting tissue-derived supernatant was titrated in a plaque assay in ST cells. Prior to homogenization, all tissues were weighed, with 0.25 g for trachea, cheek, and eyelid tissues, and 0.5 g for lung tissue. The supernatant was clarified through low-speed centrifugation. The trachea was dissected into three sections: top, middle, and bottom. The final value was calculated as the average (mean) of the titers generated from each section.

### Serological assays

96-well Maxisorp ELISA plates (Biolegend, UK) were coated overnight at 4°C, with pre-optimized concentrations of ISU-1 full-length Spike (S) protein (1 µg/ml). Two-fold dilutions of serum, BAL or nasal swab, starting from a suitable dilution in PBS-Tween plus Marvel (4%) were applied. Samples were collected either pre-inoculation (day 0) for serum only or at post-mortem. Binding of antibodies was detected with goat anti-pig IgG (Fc-specific) (AA141P, Bio-Rad, Watford, UK) or goat anti-pig IgA (AA140P, Bio-Rad, Watford, UK), conjugated to horseradish peroxidase at optimal dilutions. 1-Step™ Ultra TMB-ELISA Substrate Solution (Thermo Scientific Pierce) was added, and optical density values were read for each well at dual wavelengths (450 nm and 630 nm) using an ELx808™ Absorbance Microplate Reader (Agilent Technologies). Quantities of antibodies were determined as the reciprocal value of the dilution that yielded the first reading above the cut-off value (endpoint titer). Cut-off values were established as mean blank ODs plus 2-fold standard deviations.

### Virus neutralization assay

Virus neutralizing antibodies against all four strains of PRCV and TGEV involved were determined in serum and BAL fluid as previously described (1). Briefly 10^3^ PFU of the respective virus was incubated with an equal volume of a two-fold dilution series of serum, starting at 1:5 dilution for serum from 135-infected animals and 1:2 for serum from ISU-1-infected animals. After 30 minutes incubation at room temperature the virus/serum mix was transferred to ST cells. After 48 hours incubation at 37°C, cells were fixed and stained using 3.3% formaldehyde and 0.1% crystal violet. The presence or absence of CPE was noted, with the absence of CPE indicating neutralisation. Neutralization titers were calculated as the quantity required to achieve 50% neutralization per ml (NT50/ml) using a Reed and Muench end point calculation [48].

### ELISPot assay

MultiScreen™-HA ELISPOT plates (Millipore, Watford, UK) were coated overnight at 4°C with 0.5 µg/ml mouse anti-pig IFNγ (BD Biosciences, San Jose, CA) in carbonate buffer (0.1 M Na2CO3/NaHCO3, pH 9.6). Following three washes with PBS, the plates were incubated for two hours at 37°C with 200 µl per well of blocking buffer (RPMI, 10% FBS, supplemented with 100 U/ml penicillin and 100 µg/ml streptomycin). After removing the blocking buffer, stimuli and cells were plated. Cryopreserved cells (TBLN, BAL, spleen) were thawed and 100 µl/well at 2 x 10^6^/ml cells were added to the ELISPOT plates and cultured for 48 h at 37°C, 5% CO2 in triplicate for each condition. Stimuli included medium alone (RPMI- 1640 medium with glutamax-I and 25 mM Hepes, supplemented with penicillin, streptomycin, and 10% heat-inactivated FCS), live PRCV viruses 135 and ISU-1 (MOI 1), peptide pools covering the spike (S) of PRCV 135 virus, or 2.5 µg/mL Concanavalin A (Con A, Invitrogen). 16-mer peptides overlapping by 12 residues (Mimotopes, Melbourne, Australia) with three pools for the S protein (residues 1-100, 101-200, and 201-305) at a final concentration of 2 µg/ml were used.

After washing of the plates with PBS containing 0.05% Tween 20, 100 µl per well biotinylated mouse anti-pig IFNγ (0.5 µg/ml, BD Biosciences) was added for 2 h at room temperature. The plates were washed five times with PBS containing 0.05% Tween 20, and 100 µl per well streptavidin conjugated to alkaline phosphatase (1/1000, Invitrogen Ltd.) was added for 1 h at room temperature. Following another five washes with PBS containing 0.05% Tween 20, the plates were developed with 100 µl per well alkaline phosphatase substrate solution (Bio-Rad laboratories, Hercules, CA) for 20 min at room temperature. After rinsing with tap water, the plates were allowed to dry overnight at room temperature before counting the dark blue-coloured immunospots using the AID ELISPOT reader (AID Autoimmun Diagnostika GmbH, Strassberg, Germany). Results were expressed as the number of IFNγ-producing cells per 10^6^ stimulated cells after subtracting the average number of spots in medium-stimulated control wells.

### Intracellular cytokine staining

To assess the production of IL-2, TNF, and IFN-γ by CD8 and CD4 T cells, intracellular cytokine staining (ICS) was performed. Cell suspensions obtained from BAL, TBLN and spleen were thawed and seeded in duplicate wells of a 96-well plate (1 x 10^6^ cells per well). The cells underwent overnight stimulation with either 135 or ISU-1 (MOI 0.5), or they were treated with media (RPMI containing stable glutamine, 100 IU/ml penicillin, 100 μg/ml streptomycin, and 10% FCS) as a negative control. Incubation occurred at 37°C with 5% CO_2_. Alternatively, some cells were stimulated for 5 hours with 2 µg/ml of 100 overlapping peptides (16-mers) spanning the first 100 residues of the PRCV 135 spike protein. Positive control wells were stimulated with a 1/500 dilution of a cocktail containing phorbol 12- myristate 13-acetate (PMA) and ionomycin (BioLegend). Brefeldin A (GolgiPlug™, BD Biosciences) was added to all wells and incubated for 4 hours. Subsequently, cells were centrifuged for 5 minutes at 400g and washed twice with PBS.

Primary antibodies (**S9 Table, section A**) were applied to the cells for 20 minutes at 4°C in the dark. The cells were then permeabilized and fixed using 100 μl of BD Cytofix/Cytoperm (BD Biosciences). Following another centrifugation at 400 x g and two PBS washes, the cells were incubated for 30 minutes at 4°C in the dark with directly conjugated anti-IFN-γ, anti-TNF, and unconjugated anti-IL2 antibodies (**S9 Table, section A**). After an additional centrifugation and two washes with PBS, the cells were treated with fluorochrome-conjugated secondary antibodies. Following two more washes and resuspension in PBS, cytokine producing CD4 and CD8 T cells were analyzed using a MACSquant Analyzer 16 (Miltenyi). Wells containing only cells and medium for each animal were considered as a negative control (unstimulated), and the frequency of cytokine-producing cells was determined by subtracting the values from unstimulated cells. Data analysis was performed using FlowJo software version 10.10.0.

### Phenotypic analysis of B and T cells by flow cytometry

Thawed cell suspensions from BAL, TBLN, and spleen were resuspended in 10 ml of medium (RPMI containing stable glutamine, 100 IU/ml penicillin, 100 μg/ml streptomycin, and 10% FCS) and centrifuged at 400g for 5 minutes. Subsequently, 2 x 10^6^ cells per sample from BAL, TBLN, and spleen were seeded into a 96 U-bottom wells plate and washed twice with PBS. A cocktail of primary antibodies for either B or Tfh/ Treg phenotyping was added to the wells **(S9 Table**, **section B and section C**, respectively) and incubated for 20 minutes at 4°C. The plates were washed twice with PBS, and secondary antibodies added, followed by incubation for 20 minutes at 4°C. In cases requiring blocking, mouse serum was added. For fixation and permeabilization, eBioscience™ Foxp3 / Transcription Factor Fixation/Permeabilization Concentrate and Diluent from Invitrogen were employed for intracellular markers. Following two PBS washes, antibodies for intracellular markers were added **(S9 Table, section B and C**) to the wells and incubated for 30 minutes at 4°C. Finally, the cells were resuspended in PBS. The samples were analysed using either the MACSquant Analyzer 16 (Miltenyi) or Cytek Aurora (Cytek Biosciences). FlowJo software was employed for the analysis of the collected data.

### Luminex assay

Cells from BAL at 1 and 5 DPI were thawed and resuspended in duplicate wells of a 96-well plate (1 x 10^6 cells per well). The cells were stimulated overnight with the homologous viruses either PRCV 135 or ISU-1 at MOI 1 or they were treated with medium (RPMI containing stable glutamine, 100 IU/ml penicillin, 100 μg/ml streptomycin, and 10% FCS) as a negative control. The supernatant was collected and used for Luminex assay. Cytokine concentrations for IL-1β, IL12p40, IL-4, IL-6, IL-8, IL-10, TNF-α, IFN-γ and IFN-α were determined by Swine Cytokine Magnetic 9-plex Panel (ProcartaPlex Multiplex Immunoassay, Thermo Fisher) performed according to the manufacturer’s recommendations.

### Separation of BAL cell populations for scRNA-seq

BAL cells were isolated and subsequently cryopreserved as previously described [18, 20]. To account for the strong domination of macrophages in BAL cell preparations, we sorted major leukocyte subsets as described in [18]. In brief, thawed BAL cells were labelled with antibodies against CD4 (clone 74-12-4, PerCP-Cy5.5-conjugated, BD Biosciences), CD8β (clone PPT23, FITC-conjugated, Bio-Rad), CD3 (clone PPT3, APC-conjugated, Southern Biotech), CD16 (clone G7, AF647- conjugated, Bio-Rad), CD172a (clone 74-22-15, PE-conjugated, Bio-Rad) and Fixable Viability Dye eFluor780 (ThermoFisher) and sorted on a FACSAria UIII (BD Biosciences). This allowed the sorting of CD4^+^, CD8β^+^, CD16^+^CD172a^+^ and CD3^-^CD16^-^ cells (the latter phenotype for enrichment of B cells). Per phenotype, 12,000 cells were collected. Cells from each fraction were processed for partitioning and barcoding using a 10x Genomics Chromium iX Controller.

### Library preparation and sequencing

Single cells were processed using the Chromium iX Controller with the Next GEM chip G and the single cell 3’ kit v3.1 library preparation kit from 10x Genomics (Pleasanton, CA). For each of the BAL samples, approximately 10,000 cells were targeted for recovery during partitioning. The samples were grouped randomly and run in pairs, using three NextSeq High Output 150 cycle reagent kits (Illumina, San Diego, CA), with 1% PhiX, aiming for about 200 million reads per sample, with the run format aligning with 10X guidelines. The sequencing was performed on an Illumina NextSeq 550, using 3 high output 200 cycle reagent kit and flow cells. Bcl files were demultiplexed using ‘cellranger mkfastq’ and aligned/counted with ‘cellranger count’ (cellranger-7.0.0), using default parameters and the Sus scrofa genome (genome assembly 11.1, Ensembl release 108).

### scRNA-seq QC and pre-processing

Data analysis was conducted using R (version 4.3.2). To ensure data integrity, we excluded empty droplets and barcode-swapped droplets from the unfiltered CellRanger output using DropletUtils (1.22.0) [49, 50]. Further filtering involved the removal of cells with total UMI counts exceeding 50,000 or falling below 500. Genes without any counts and cells exhibiting an outlying proportion of mitochondrial UMIs (greater than 5 median absolute deviations from the median) were also excluded.

Normalization was achieved using pooled factors [51], and samples from the same sequencing run were merged to form a single analysis dataset. Specifically, samples 1-8 from sequencing run 1 were consolidated to create the 1 DPI analysis dataset, while samples 9-16 from sequencing run 2 were combined to form the 20 DPI analysis dataset. Batch effect correction was deemed unnecessary as visualisation and clustering revealed no batch effects within each run.

### scRNA-seq clustering and annotation

Shared-nearest-neighbors graph-based clustering was performed on each analysis dataset using clusterCells (bluster 1.12.0) with the top 5000 highly variable genes (HVGs) identified by batchelor (1.18.1) [52]. The clustering was conducted using the Jaccard index to weight edges and the Louvain method for community detection, with a k value of 20.

For the 1 DPI dataset, the clustering process generated 19 initial clusters. Cluster 18, which exhibited particularly low counts, had a limited number of genes detected, and showed dispersive distribution in UMAP. Therefore, it was considered as poor-quality and excluded from further analysis. Cluster 14 in the 1 DPI dataset displayed a dichotomy of CD4 and CD8 expression. This cluster was further split into CD4, CD8, and NK subclusters. The sub-clustering process involved identifying the top 1000 HVGs within the cluster, selecting those genes that correlated best with CD4 and CD8B expression using a zero-inflated Kendall’s Tau correlation metric implemented in scHOT (v1.14.0), and re-running the clustering command using Jaccard index to weight edges and the fast greedy modularity optimization algorithm for community detection on cluster 14 based on only those HVGs. Similarly, in the 20 DPI dataset, cluster 9 exhibited a dichotomy of CD4 and CD8 expression. This cluster was split into CD4+ and CD8β+ subclusters using the same strategy as described for cluster 14 in the 1 DPI dataset.

In the 1 DPI dataset, varied expression of genes including KLRB1 and NCR1 within Cluster 9 prompted sub-clustering. This was performed in an unsupervised manner by identifying the top 2000 HVGs in cluster 9 cells and then re-running the clustering using these HVGs. The Jaccard index was used to weight edges, and the fast greedy modularity optimization algorithm was applied for community detection, resulting in the division of Cluster 9 into two subclusters. Using the same approach, cluster 12 was split into three subclusters, and Cluster 3 into two subclusters.

The 20 DPI dataset generated 20 initial clusters. In this dataset, clusters 2, 4, 15, and 19 were each split into two subclusters using the unsupervised method described above to better distinguish and define cell types within. Subclusters containing fewer than 100 cells were excluded from further analysis due to their inability to produce significant statistical results.

Clusters were manually annotated with conventional names (cell types) by examining expression of typical defining markers, as listed in **S1 Table** and illustrated in **Figure 5A** and **Supl Fig S4**. Additionally, top marker genes within each cluster were determined using the score Markers function (scran 1.30.2), as detailed in **S10 and S11 Tables**.

### scRNA-seq analysis, differential abundance analyses

To compare the cell type proportion (abundance of cells within each cluster) across two virus infection conditions, we performed differential abundance analyses using a negative binomial generalized linear model with empirical Bayes quasi-likelihood F-tests [53, 54], as implemented in edgeR (v4.0.16). The model was fitted with two conditions (135 vs. ISU-1) to test for differential abundance in condition 135, using ISU-1 as the baseline. Trend estimation was disabled when estimating dispersion and quasi-likelihood dispersion.

To account for the sort and pool strategy used to balance cell types among the sequenced cells, clusters were divided according to their membership in the four sorted cell populations (macrophages, CD4 T cells, CD8 T cells, B cells) before running differential abundance analyses within each group (in **S2 Table**). Clusters that did not clearly fall into one of these cell types were excluded to ensure the universes for differential abundance analysis were equivalent to or smaller than the sorted cell populations. This approach was intended to avoid sorting biases in intra-cell-type differential abundance analyses.

### scRNA-seq analysis, differential expression analyses

Due to the limited number of subjects in each condition we conducted pseudo-bulk differential expression analyses within each cluster using edgeR (**Figure 6A, 6B, 7A**, **Sup Fig S7-8, S10**). To compare between 135 and ISU-1 infections, counts from all cells within each cluster were pooled for each sample and clusters with fewer than 10 cells within each sample were removed. A negative binomial generalized linear model was then fitted for each cluster and differential expression between conditions was tested using empirical Bayes quasi-likelihood F-tests, as implemented in edgeR (v4.0.16). The differential expression results are provided in **S12 and S13 Tables.**

Afterwards, we first combined all genes that were differentially expressed in any cluster within each major cell population (e.g. CD4 T cells) (**S5 Table**). Then, we created Venn diagrams to display those DE genes within four main cell populations (macrophages, CD4 T cells, CD8 T cells, and B cells). We also checked the direction of effect in cluster within each main population; these directions were all consistent across clusters from the same main population.

### Gene Ontology Enrichment

Lists of differentially expressed genes were analyzed for Gene Ontology (GO) enrichment using clusterProfiler (4.10.1), with the genome-wide annotation for pig (DOI: 10.18129/B9.bioc.org.Ss.eg.db, release 3.19), focusing on the biological process sub-ontology.

When using clusterProfiler, hypergeometric testing was employed to identify GO terms significantly enriched in the set of DE genes in each cluster compared to a background set of all measured genes. To account for multiple testing, the Benjamini-Hochberg (BH) method was applied to adjust p-values. GO terms with an adjusted p-value (q-value) < 0.05 were considered statistically significant and are displayed in bubble plots. All results are available in **S7 and S8 Tables**. Since results of 1 DPI only contained very few significant GO terms in per cell type, no bubble plots were created.

### Statistical analysis

GraphPad Prism 9.2.0 (GraphPad Software, San Diego, CA, United States) was used for statistical analyses. The specific type of analysis used is indicated in each figure legend. Data were first tested for normality. For normally distributed data sets a one-way or two-way ANOVA was applied followed by Bonferroni’s multiple comparisons test. For non-normally distributed data a non-parametric unpaired t test was used. A p-value of <0.05 was considered to be statistically significant. To assess correlation between virological/immune parameters and pathological measures, a non-parametric Spearman correlation coefficient (*ρ*) was computed for each pair of measures (viral load, antibody response, T cell responses and pathology) using Graphpad Prism v 10.0.3. All 135- and ISU-1- infected samples at the specified time points were included in the analysis.

## Supplementary Material

**S1 Table. scRNA-seq cluster and cell type annotation**. Table showing all clusters passing quality control, their annotated cell type, and the list of HGNC genes used to delineate cell types.

**S2 Table. Cluster membership in the four sorted cell populations in differential abundance analysis.**

**S3 Table**. **Differential abundance.** Results from differential abundance analyses between two conditions using a negative binomial generalized linear model with empirical Bayes quasi-likelihood F-tests. Including tests for each cluster identified to belong to any of the sorted cell populations: CD4 T cells (CD4), CD8 T cells (CD8), monocytes, macrophages, dendritic cells (MMD) or B cells and plasma cells (B).

**S4 Table. Shared differentially expressed genes between cell types 1 DPI.** HGNC gene names suffixed with ‘-Up’ imply significant upregulation in that cell type(s) in 135 condition, HGNC gene names suffixed with ‘-Down’ imply significant downregulation in that cell type(s) in 135 infected samples at 1 DPI.

**S5 Table. Cluster assignment.** Clusters assigned to each main cell population in Venn diagrams, shared DE genes tables and GO bubble plots.

**S6 Table. Differentially expressed shared genes between cell types 20 DPI**. HGNC gene names suffixed with ‘-Up’ imply significant upregulation in that cell type(s) in 135 condition, HGNC gene names suffixed with ‘-Down’ imply significant downregulation in that cell type(s) in 135 infection at 20 DPI.

**S7 Table. GO enrichment of terms shared between B cells, CD4 and CD8 T cells (20 DPI).** GO enrichment results using clusterProfiler package, showing the significant GO terms in 135 infection as tested against ISU-1 shared between B cells, CD4 and CD8 T cells.

**S8 Table. GO enrichment of terms for all cell types (1 DPI and 20 DPI).** GO enrichment results using clusterProfiler package, showing all significant GO terms appeared within each main populations in 135 infected samples as tested against ISU-1.

**S9 Table. Primary antibodies and second-step reagents used in flow cytometry.** Details on antibodies for each staining panel (intracellular cytokine staining, B cells and plasma cells, Tfh and Tregs) are listed.

**S10 Table. Gene expression scores produced by scoreMarkers at 1DPI** The top marker genes within each cluster were identified by the scoreMarkers function (scran 1.30.2) to produce lists of genes and scores, ranking genes by their degree of distinctive expression versus all other clusters.

**S11 Table Gene expression scores produced by scoreMarkers 20 DPI.** The top marker genes within each cluster were identified by the scoreMarkers function (scran 1.30.2) to produce lists of genes and scores, ranking genes by their degree of distinctive expression versus all other clusters.

**S12 Table edgeR differential expression results 1 DPI.** The differential expression results within each cluster were identified by edgeR package (v4.0.16). The lists of genes and scores, ranking genes by adjusted p values.

**S13 Table edgeR differential expression results 20 DPI.** The differential expression results within each cluster were identified by edgeR package (v4.0.16). The lists of genes and scores, ranking genes by adjusted p values.

## Author Contributions

**Conceptualization:** Elma Tchilian, Sarah Keep, Erica Bickerton, Graham Freimanis, Wilhelm Gerner

**Data curation:** Ehsan Sedaghat-Rostami, Brigid Veronica Carr, Liu Yang, Sarah Keep, Andrew Muir, Isabella Atkinson, Albert Fones, Katy Moffat, Graham Freimanis

**Formal analysis:** Ehsan Sedaghat-Rostami, Brigid Veronica Carr, Liu Yang, Sarah Keep, Fabian Z X Lean, Basudev Paudyal, James Kirk, Simon Gubbins, Adam MacNee, Isabella Atkinson, Albert Fones, Emily Briggs, Katy Moffat, Graham Freimanis, Andrew Muir, Arianne Richard, Nicos Angelopoulos, Elma Tchilian

**Funding acquisition:** Elma Tchilian, Sarah Keep, Erica Bickerton, Graham Freimanis

**Investigation:** Ehsan Sedaghat-Rostami, Brigid Veronica Carr, Liu Yang, Sarah Keep, Fabian Z X Lean, Basudev Paudyal, Adam MacNee, Emily Briggs, Alejandro Nunez, Katy Moffat, Graham Freimanis, Andrew Muir, Arianne Richard, Nicos Angelopoulos, Wilhelm Gerner, Elma Tchilian

**Methodology:** Ehsan Sedaghat-Rostami, Brigid Veronica Carr, Liu Yang, Sarah Keep, Fabian Z X Lean, Basudev Paudyal, James Kirk, Eleni Vatzia, Simon Gubbins, Erica Bickerton, Adam MacNee, Emily Briggs, Alejandro Nunez, Katy Moffat, Graham Freimanis, Christine Rollier, Andrew Muir, Arianne Richard, Nicos Angelopoulos, Wilhelm Gerner, Elma Tchilian

**Project administration:** Elma Tchilian

**Resources:** Elma Tchilian, Arianne C. Richard, Nicos Angelopoulos

**Software:** Andrew Muir, Arianne C. Richard, Nicos Angelopoulos, Liu Yang

**Supervision:** Wilhelm Gerner, Arianne C. Richard, Nicos Angelopoulos, Elma Tchilian

**Validation:** Ehsan Sedaghat-Rostami, Andrew Muir, Nicos Angelopoulos, Arianne C. Richard, Liu Yang, Simon Gubbins, Wilhelm Gerner, Elma Tchilian

**Visualization:** Ehsan Sedaghat-Rostami, Brigid Veronica Carr, Liu Yang, Sarah Keep, Fabian Z X Lean, Basudev Paudyal, Wilhelm Gerner, Arianne C. Richard, Andrew Muir, Nicos Angelopoulos, Wilhelm Gerner, Elma Tchilian

**Writing – original draft:** Elma Tchilian, Ehsan Sedaghat-Rostami, Wilhelm Gerner, Fabian Z X Lean, Liu Yang

**Writing – review & editing:** Ehsan Sedaghat-Rostami, Brigid Veronica Carr, Liu Yang, Sarah Keep, Fabian Z X Lean, Basudev Paudyal, Simon Gubbins, Erica Bickerton, Emily Briggs, Alejandro Nunez, Katy Moffat, Graham Freimanis, Christine Rollier, Andrew Muir, Arianne C. Richard, Nicos Angelopoulos, Wilhelm Gerner, Elma Tchilian

## Acknowledgments

We thank the flow cytometry, sequencing and bioinformatics facilities at The Pirbright Institute for enabling this research. We are grateful to the staff at Animal and Plant Health Agency for providing excellent animal care.

## Supplementary figure legends

**Supplementary Figure 1.**
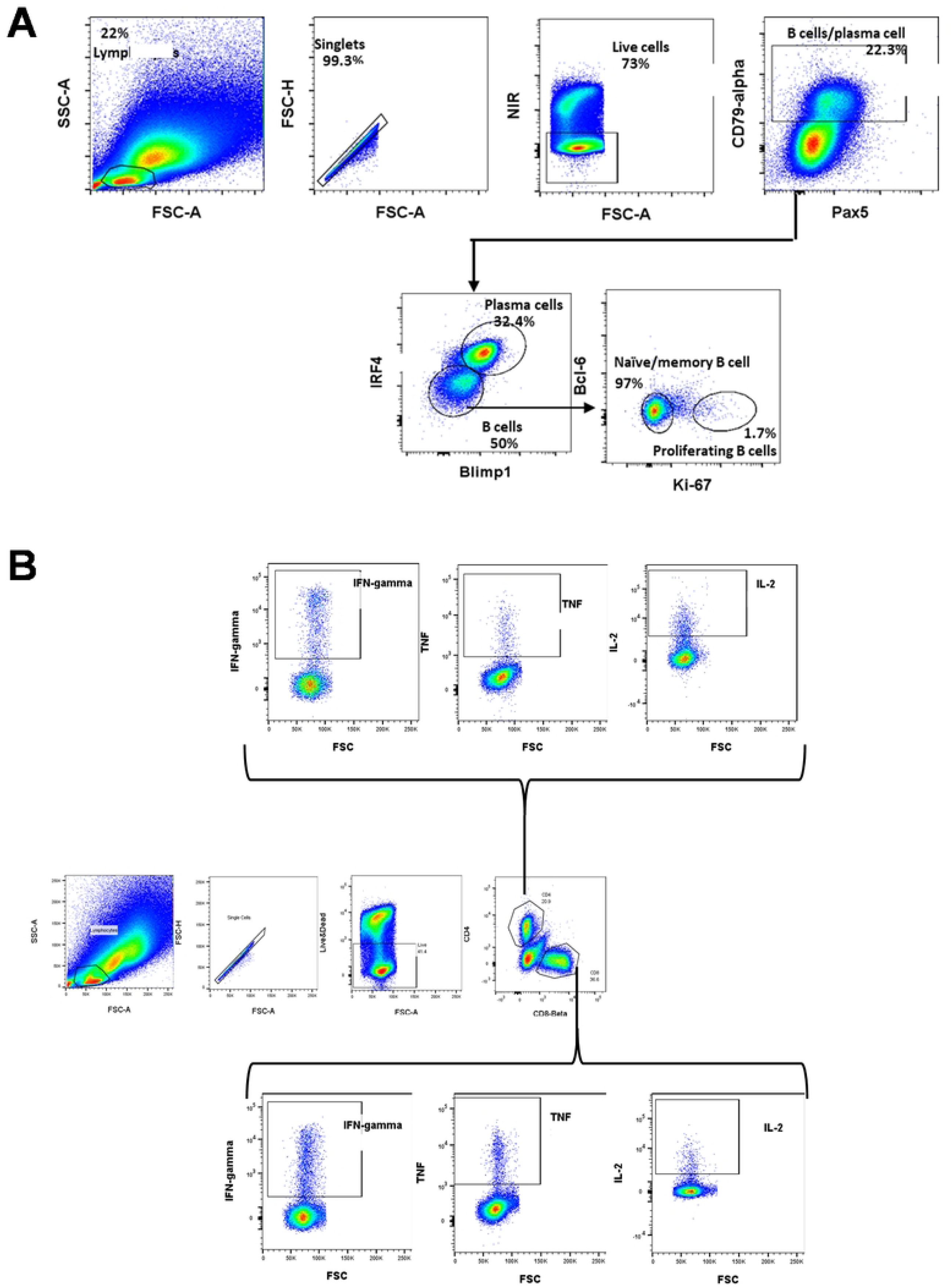
Gating strategy for B cell and intracellular cytokine staining in T cells. **(A)** Cryopreserved BAL samples of both PRCV ISU-1 and 135 were thawed and flow cytometry staining performed. Sequential gating strategy was applied (left to right) to identify the frequency B cells subsets. **(B)** Cryopreserved BAL samples were thawed and stimulated with PMA/I, 135, ISU-1, Spike peptide pools or medium. Flow cytometry staining was performed. Sequential gating strategy was applied on lymphocytes, singlets, live cells and CD4 or CD8-beta. The frequencies of IFN-γ, TNF and IL-2 secreting CD4 and CD8 T cells also were analyzed.

**Supplementary Figure 2.**
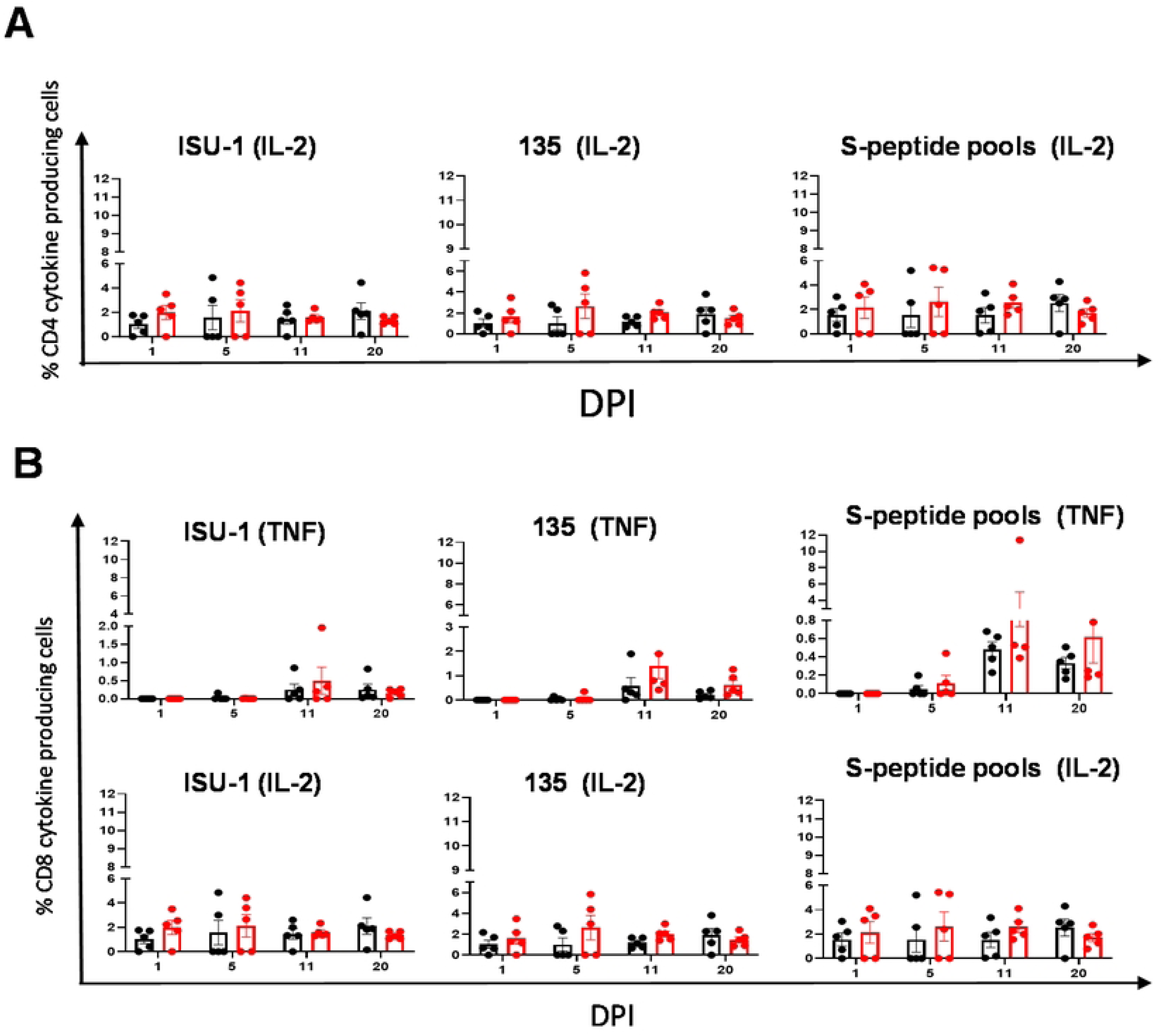
Cytokine T cell responses in BAL following ISU-1 or 135 infections. **(A)** The frequency of IL-2 secreting CD4 and **(B)** TNF and IL-2 secreting CD8 T cells were determined using intracellular staining following stimulation with ISU-1, 135 or peptide pools covering the Spike protein. Each point represents single animal. The line on each bar represent mean with SEM. Each point represents single animal and mid-line in the middle represent the mean. Two-way ANOVA and Bonferroni’s multiple test were performed. Asterisks represent significant changes (*p < 0.05, **p < 0.01, ***p < 0.001, ****p < 0.0001).

**Supplementary Figure 3.**
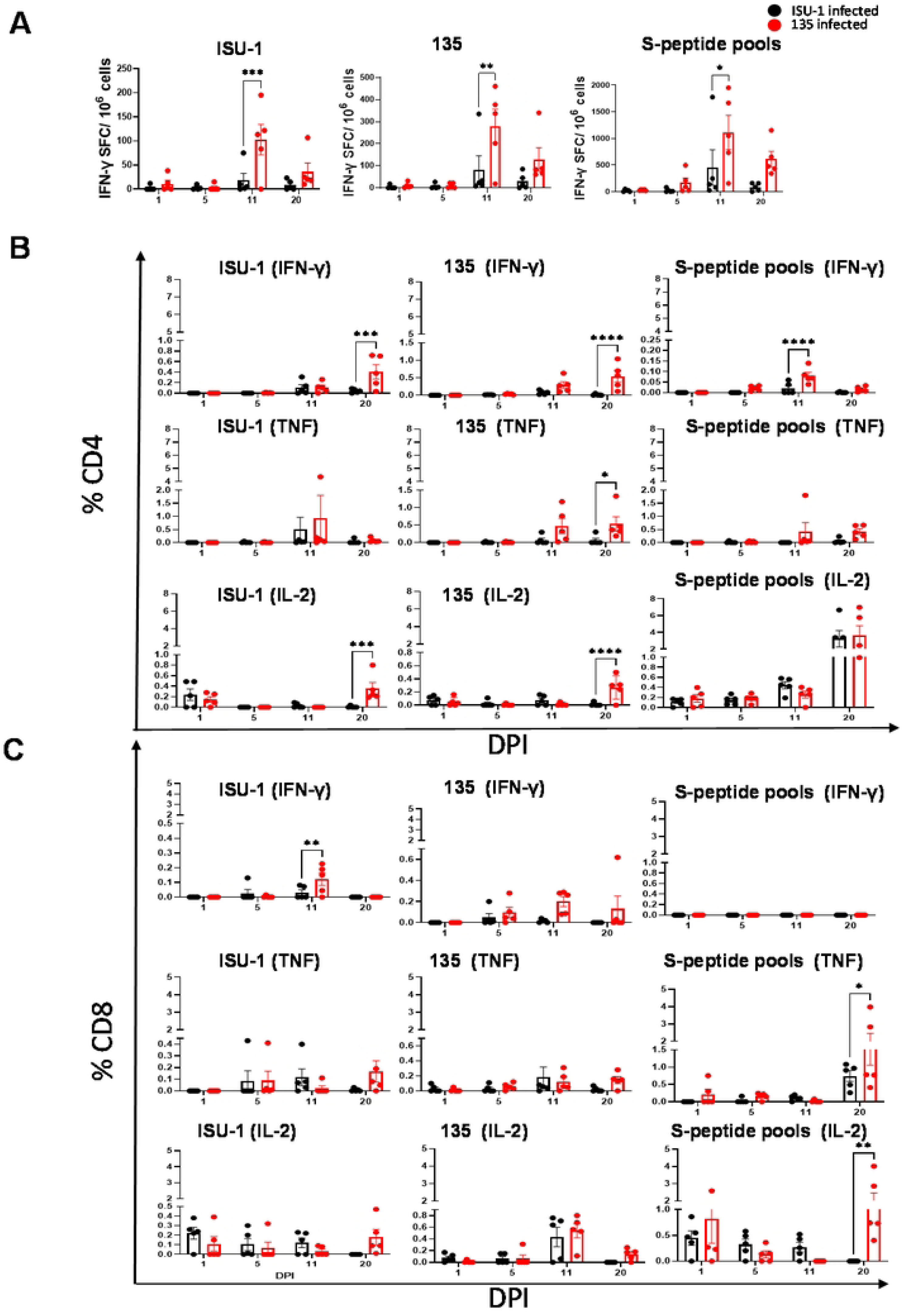
Cytokine T cell responses in TBLN following ISU-1 or 135 infections. **(A)** IFN-γ secreting spot forming cells (SFC) were enumerated by ELISpot following *ex-vivo* stimulation with ISU-1, 135 or spike peptide pools. The frequency of IFN-γ, TNF and IL-2 secreting CD4 **(B)** and CD8 **(C)** T cells were determined using intracellular staining following stimulation with ISU-1, 135 or peptide pools covering the Spike protein. Each point represents single animal. The line on each bar represent mean with SEM. Each point represents single animal and mid-line in the middle represent the mean. Two-way ANOVA and Bonferroni’s multiple test were performed. Asterisks represent significant changes (*p < 0.05, **p < 0.01, ***p < 0.001, ****p < 0.0001).

**Supplementary Figure 4.**
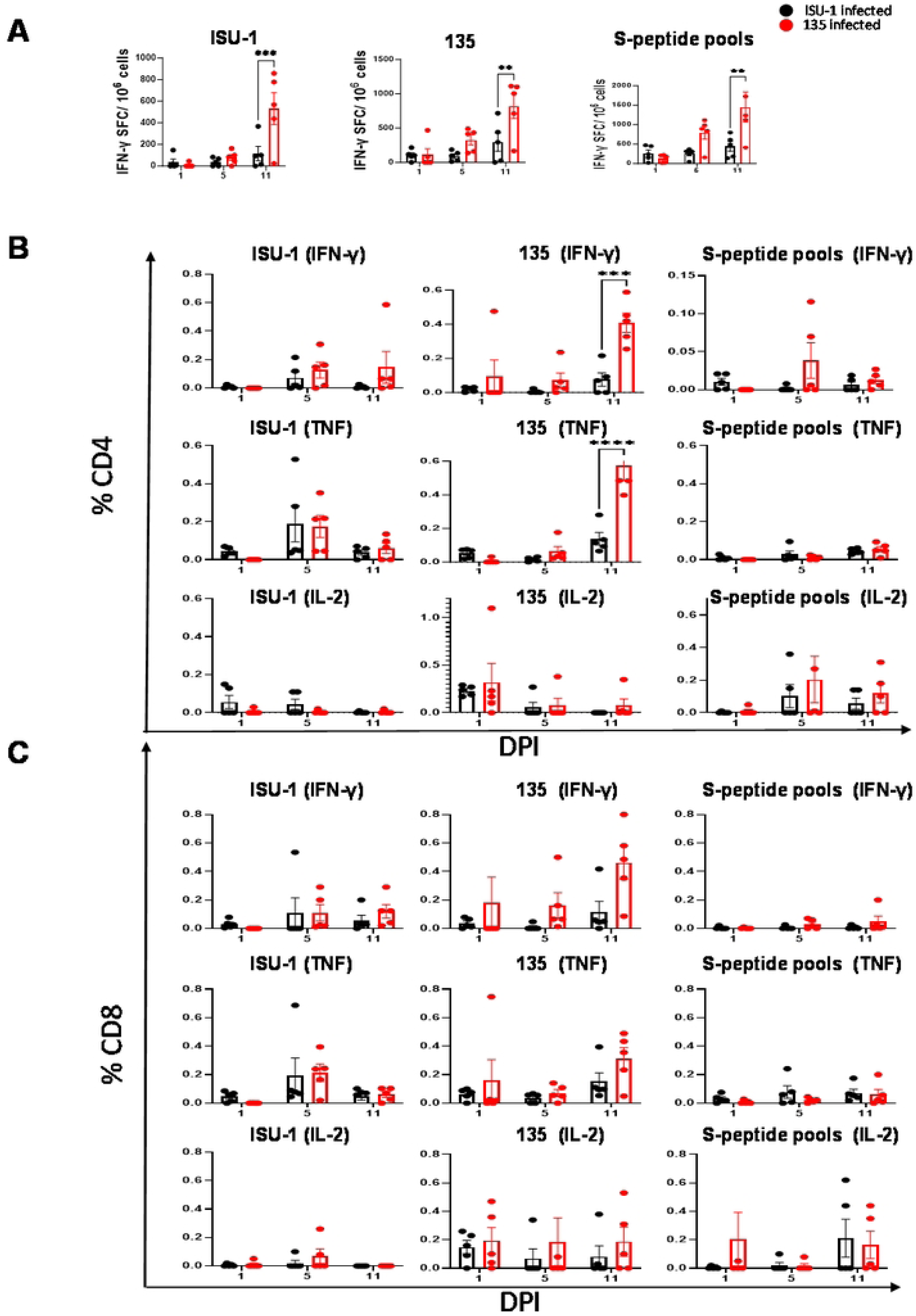
Cytokine T cell responses in spleen following ISU-1 or 135 infections. **(A)** IFN-γ secreting spot forming cells (SFC) were enumerated by ELISpot following *ex-vivo* stimulation with ISU-1, 135 or spike peptide pools. The frequency of IFN-γ, TNF and IL-2 secreting CD4 **(B)** and CD8 **(C)** T cells were determined using intracellular staining following stimulation with ISU-1, 135 or peptide pools covering the Spike protein. Each point represents single animal. The line on each bar represent mean with SEM. Each point represents single animal and mid-line in the middle represent the mean. Two-way ANOVA and Bonferroni’s multiple test were performed. Asterisks represent significant changes (*p < 0.05, **p < 0.01, ***p < 0.001, ****p < 0.0001).

**Supplementary Figure 5.**
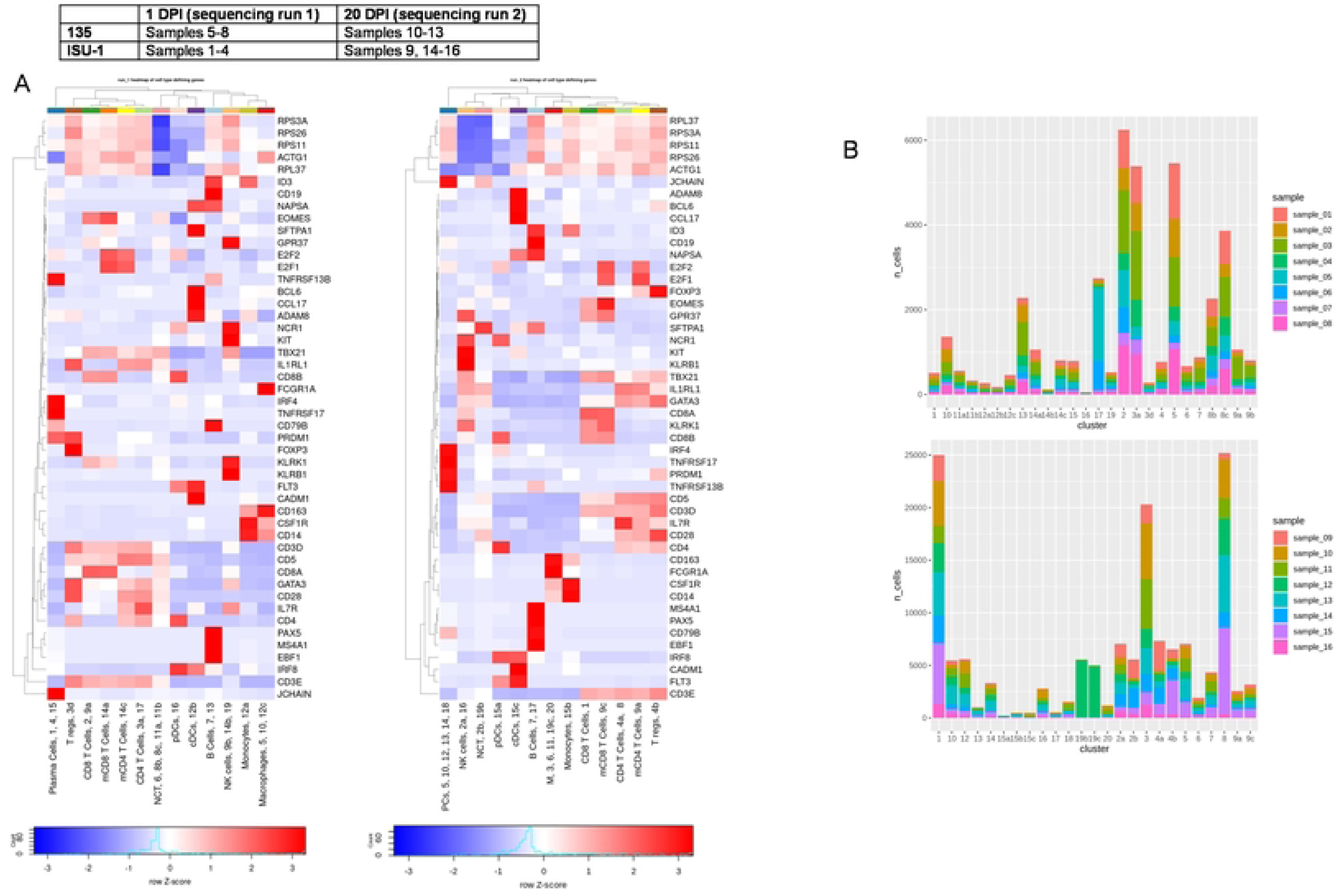
**(A)** Heatmaps of canonical marker gene expression across all scRNA-seq clusters. Left: 1 DPI, right 20 DPI. non-conventional T cells (NCT), Macrophages (M), Mitotic CD8 T Cells (mCD8 T Cells). **(B)** Sample contribution to each cluster. Top: 1 DPI. Bottom: 20 DPI.

**Supplementary Figure 6.**
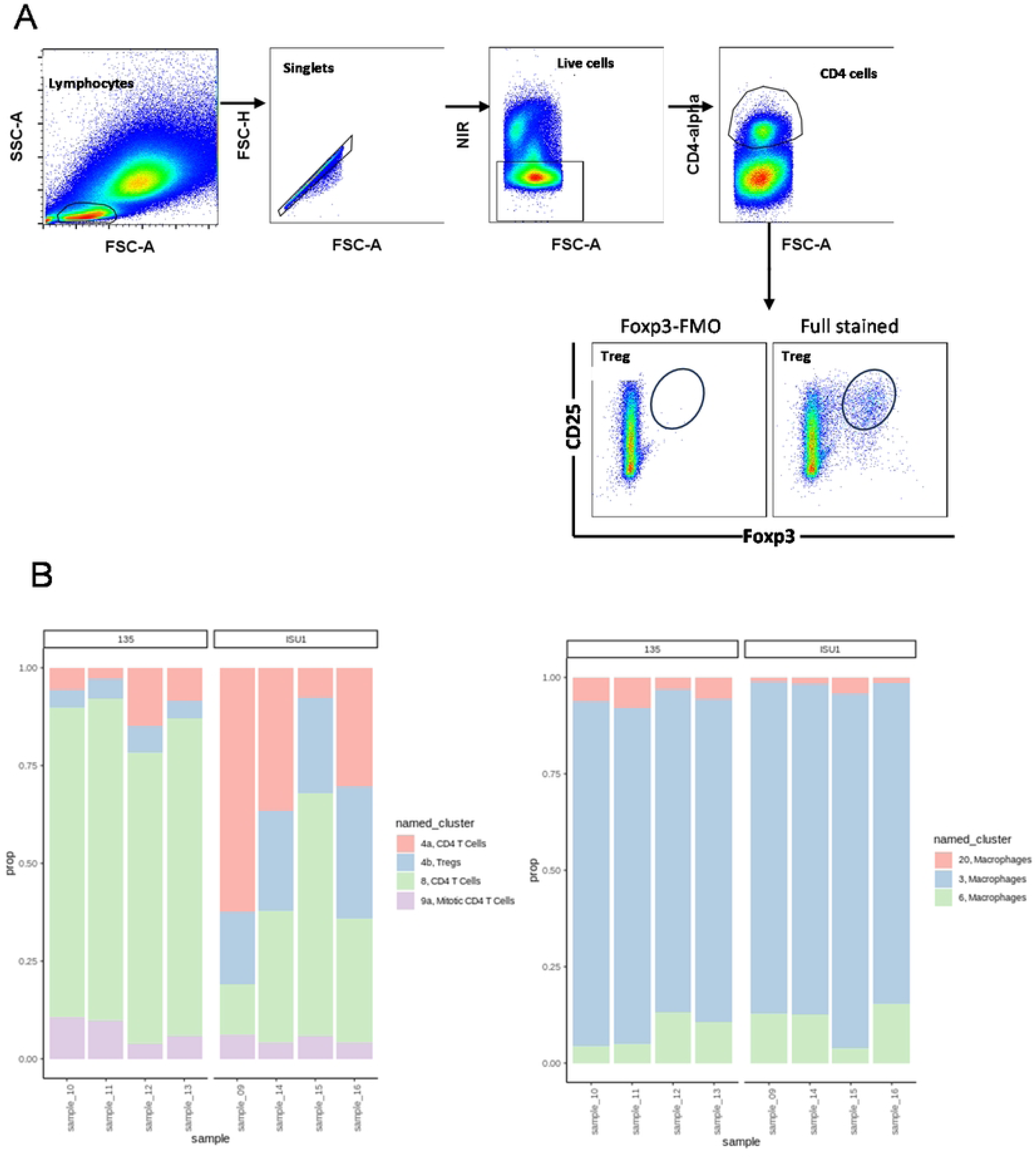
Gating strategy for Treg in BAL and composition of CD4 T cell/ macrophage clusters by scRNA-seq. (A) BAL samples were thawed, and flow cytometry staining performed. Sequential gating was applied to identify the frequency of T regulatory cells, based on CD4, CD25 and Foxp3 expression. (B) left: Proportions of clusters 4a, 4b, 8, and 9a in sorted CD4 T cell population across all BAL samples at 20 DPI by scRNA-seq. Right: Proportions of clusters 3, 6, 11 and 20 in sorted macrophage population across all BAL samples at 20 DPI. Virus strains are indicated at the top of the graphs.

**Supplementary Figure 7.**
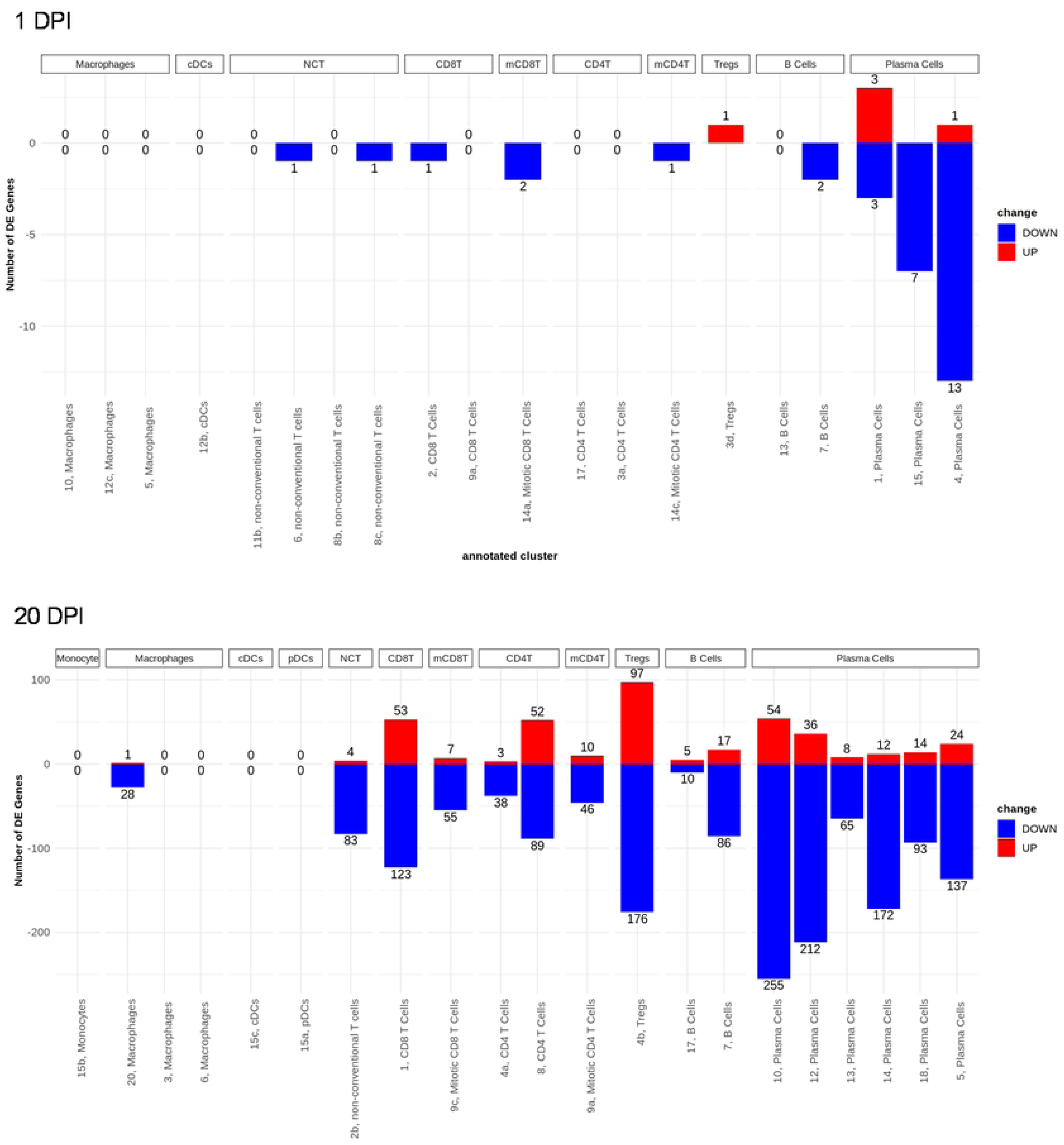
Overview of the number of DE genes between 135- and ISU-1- infected pigs within each cluster. Top: 1 DPI, bottom: 20 DPI. Each column illustrates the numbers of genes with higher or lower expression in 135-infected pigs per cluster; with cells from ISU-1-infected pigs serving as the reference.

**Supplementary Figure 8.**
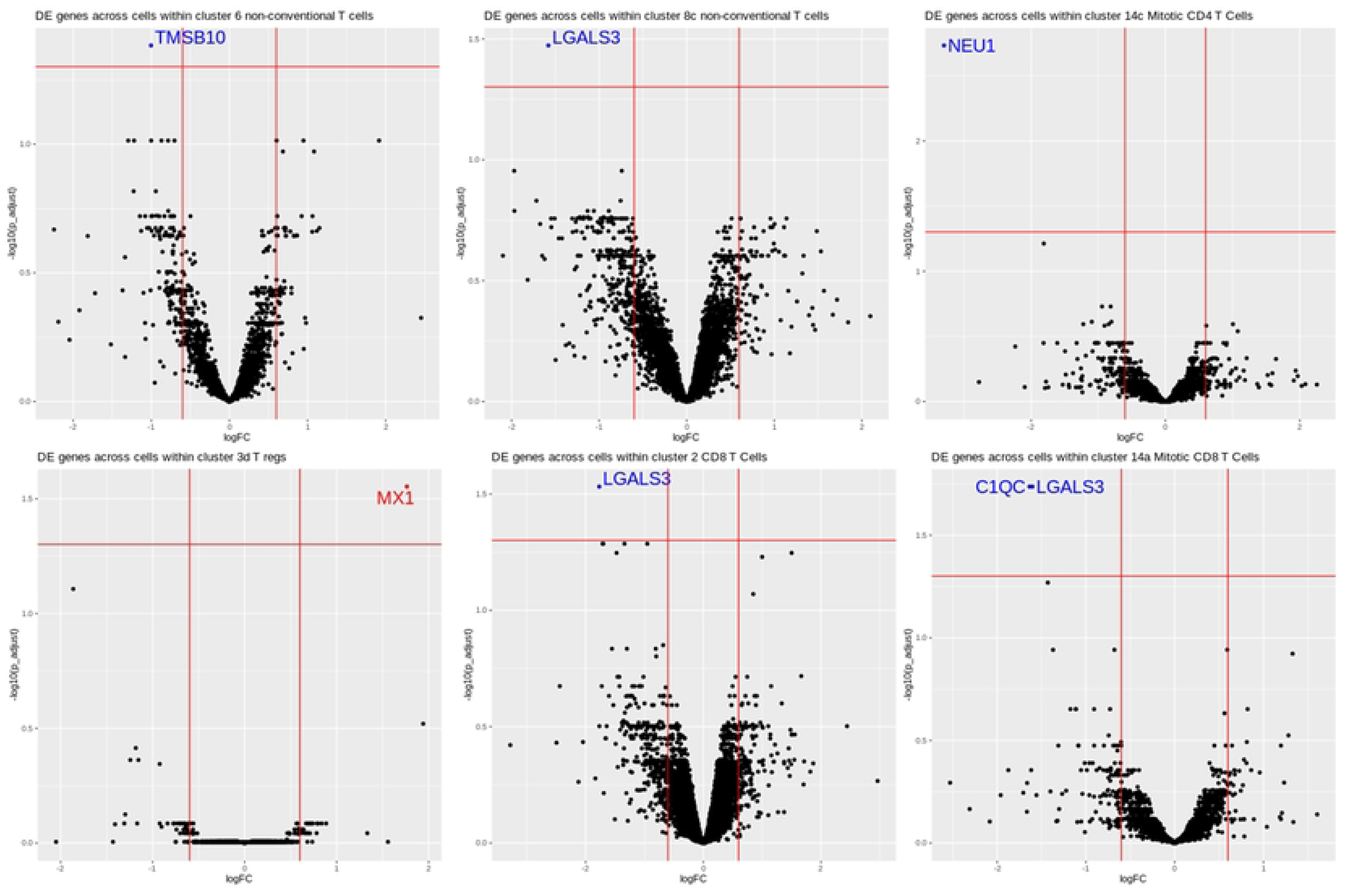
Volcano plots of DE genes between 135- and ISU-1-infected pigs within each cluster at 1 DPI. Each plot represents one cluster. Cells from ISU1-infected pigs serve as the reference. Horizontal red line is at –log10(1.25). Vertical red lines are at −0.6 and 0.6 logFC. Adjusted p-values were calculated with the Benjamini Hochberg procedure.

**Supplementary Figure 9.**
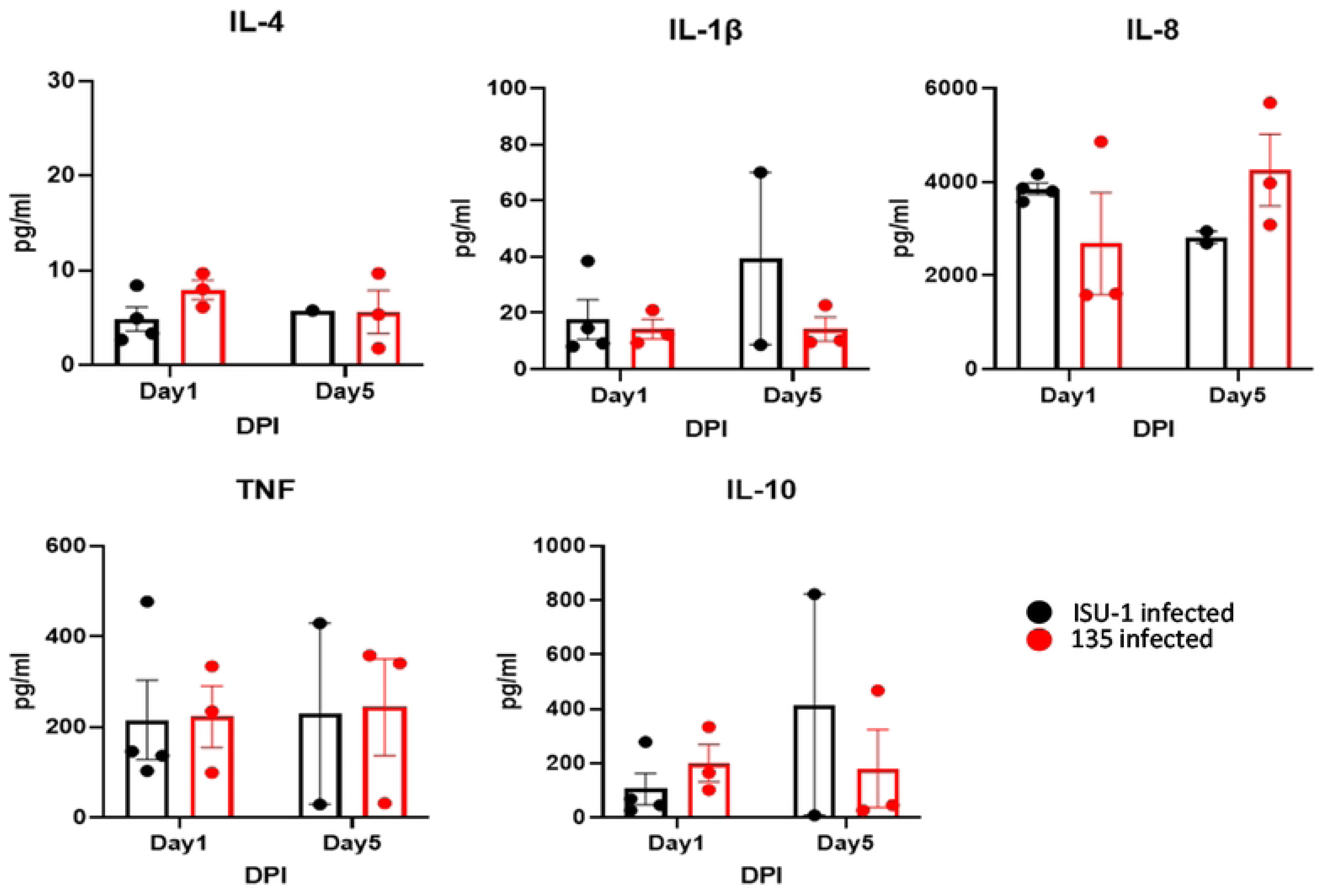
Cytokines in BAL following ISU-1 or 135 infections. BAL cells from ISU-1 (black) and 135 (red) infected pigs were stimulated overnight with the respective homologous virus strain. The supernatant was collected and Luminex assay was performed to measure the concentration of IL-4, IL-1β, IL-8, TNF and IL-10. Each dot represents a single animal.

**Supplementary Figure 10.**
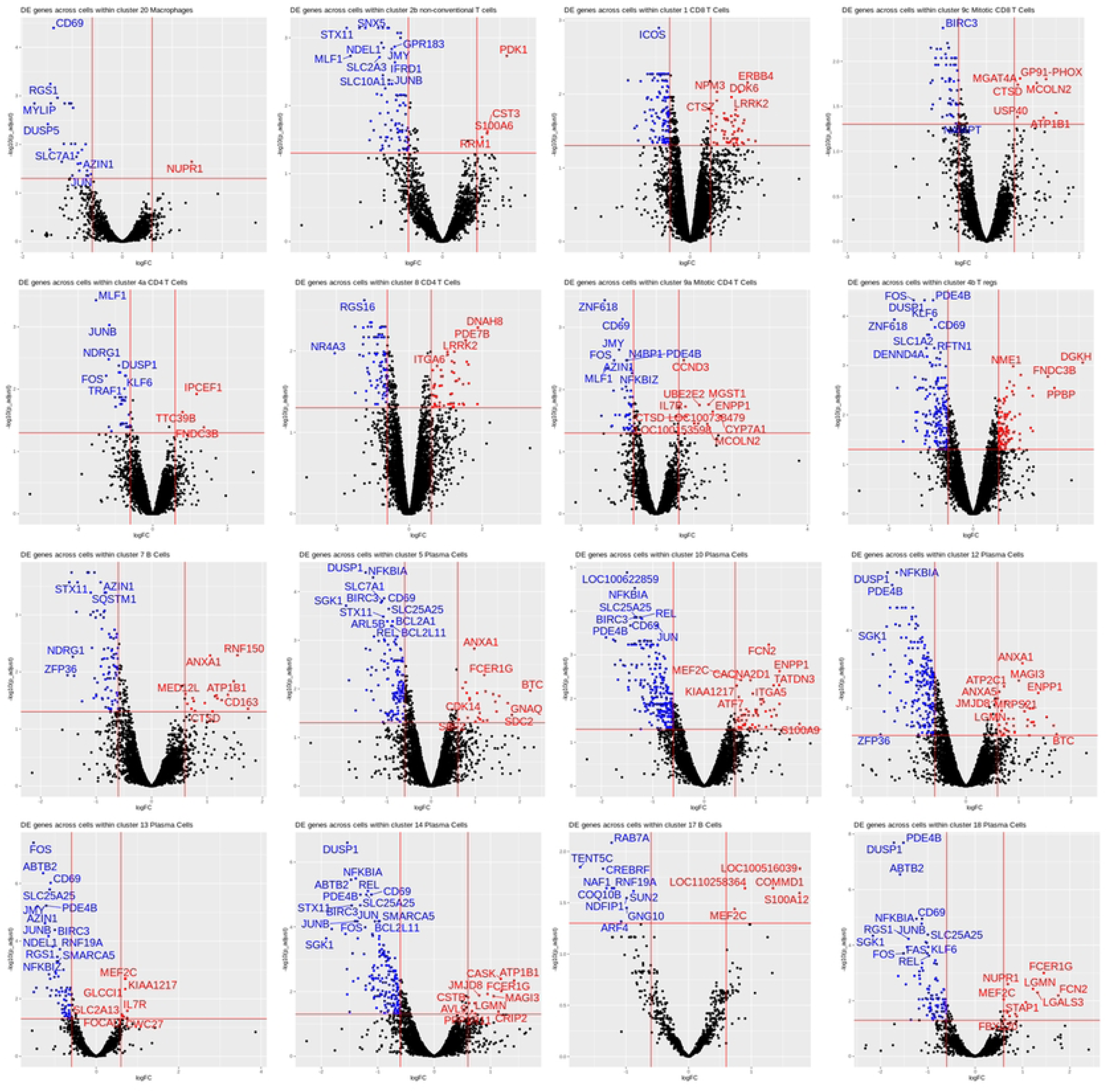
Volcano plots of DE analysis results between 135- and ISU- 1-infected pigs within each cluster at 20 DPI. Each plot displays DE analysis results within one cluster. Cells from ISU1-infected pigs serve as the reference. Horizontal red line is at – log10(1.25). Vertical red lines are at −0.6 and 0.6 logFC. Adjusted p-values were calculated with the Benjamini Hochberg procedure.

**Supplementary Figure 11. GO terms identified in B cells, CD4 T cells or CD8 T cells at 20 DPI.** Enriched GO terms, based on all genes with an adjusted p value of less than 0.05 for either B cells, CD4 T cells or CD8 T cells, or shared between these cell types, are shown. Bubble size represents the ratio of the number of DE genes associated with a specific GO term to the total number of DE genes identified in that cluster. Bubble colour represents the adjusted p-value. Terms are ranked according to gene ratio values. Macrophage populations did not show significantly enriched GO terms.

## References

1. Saif, L.J. and K. Jung, Comparative Pathogenesis of Bovine and Porcine Respiratory Coronaviruses in the Animal Host Species and SARS-CoV-2 in Humans. J Clin Microbiol, 2020. 58(8).

2. Khamassi Khbou, M., et al., Coronaviruses in farm animals: Epidemiology and public health implications. Vet Med Sci, 2021. 7(2): p. 322–347.

3. Pensaert, M., P. Callebaut, and J. Vergote, Isolation of a porcine respiratory, non-enteric coronavirus related to transmissible gastroenteritis. Vet Q, 1986. 8(3): p. 257–61.

4. Rasschaert, D., M. Duarte, and H. Laude, Porcine respiratory coronavirus differs from transmissible gastroenteritis virus by a few genomic deletions. J Gen Virol, 1990. 71 **(Pt** **11**): p. 2599–607.

5. Saif, L.J., J.L. van Cott, and T.A. Brim, Immunity to transmissible gastroenteritis virus and porcine respiratory coronavirus infections in swine. Vet Immunol Immunopathol, 1994. 43(1- 3): p. 89–97.

6. Vlasova, A.N., et al., Porcine coronaviruses. . Emerging and Transboundary Animal Viruses, 2020: p. 79–110.

7. Jung, K., et al., Altered pathogenesis of porcine respiratory coronavirus in pigs due to immunosuppressive effects of dexamethasone: implications for corticosteroid use in treatment of severe acute respiratory syndrome coronavirus. J Virol, 2007. 81(24): p. 13681–93.

8. Keep, S., et al., Porcine respiratory coronavirus as a model for acute respiratory coronavirus disease. Front Immunol, 2022. 13:867707.

9. Delmas, B., et al., Further Characterization of Aminopeptidase-N as a Receptor for Coronaviruses, in Coronaviruses: Molecular Biology and Virus-Host Interactions, H. Laude and J.-F. Vautherot, Editors. 1993, Springer US: Boston, MA. p. 293–298.

10. Delmas, B., et al., Determinants essential for the transmissible gastroenteritis virus-receptor interaction reside within a domain of aminopeptidase-N that is distinct from the enzymatic site. J Virol, 1994. 68(8): p. 5216–24.

11. Peng, J.Y., et al., Time-dependent viral interference between influenza virus and coronavirus in the infection of differentiated porcine airway epithelial cells. Virulence, 2021. 12(1): p. 1111–1121.

12. Paul, P.S., E.M. Vaughn, and P.G. Halbur, Pathogenicity and sequence analysis studies suggest potential role of gene 3 in virulence of swine enteric and respiratory coronaviruses. Adv Exp Med Biol, 1997. 412: p. 317–21.

13. Muñoz-Fontela, C., et al., Animal models for COVID-19. Nature, 2020. 586(7830): p. 509–515.

14. Flerlage, T., et al., Influenza virus and SARS-CoV-2: pathogenesis and host responses in the respiratory tract. Nat Rev Microbiol, 2021. 19(7): p. 425–441.

15. Salguero, F.J., et al., Comparison of rhesus and cynomolgus macaques as an infection model for COVID-19. Nat Commun, 2021. 12(1): p. 1260.

16. Pabst, R., The pig as a model for immunology research. Cell Tissue Res, 2020. 380(2): p. 287–304.

17. Judge, E.P., et al., Anatomy and bronchoscopy of the porcine lung. A model for translational respiratory medicine. Am J Respir Cell Mol Biol, 2014. 51(3): p. 334–43.

18. Muir, A., et al., Single-cell analysis reveals lasting immunological consequences of influenza infection and respiratory immunization in the pig lung. PLoS Pathog, 2024. 20(7): p. e1011910.

19. Nasiri, N., et al., Ocular Manifestations of COVID-19: A Systematic Review and Meta-analysis. J Ophthalmic Vis Res, 2021. 16(1): p. 103–112.

20. Schmidt, A., et al., Effect of mucosal adjuvant IL-1β on heterotypic immunity in a pig influenza model. Front Immunol, 2023. 14: p. 1181716.

21. Son, Y.M., et al., Tissue-resident CD4(+) T helper cells assist the development of protective respiratory B and CD8(+) T cell memory responses. Sci Immunol, 2021. 6(55).

22. Katoh, S., et al., A crucial role of sialidase Neu1 in hyaluronan receptor function of CD44 in T helper type 2-mediated airway inflammation of murine acute asthmatic model. Clin Exp Immunol, 2010. 161(2): p. 233–41.

23. Chen, H.Y., et al., Galectin-3 negatively regulates TCR-mediated CD4+ T-cell activation at the immunological synapse. Proc Natl Acad Sci U S A, 2009. 106(34): p. 14496–501.

24. Wagner, E.F. and R. Eferl, Fos/AP-1 proteins in bone and the immune system. Immunol Rev, 2005. 208: p. 126–40.

25. Tomasello, E. and E. Vivier, KARAP/DAP12/TYROBP: three names and a multiplicity of biological functions. Eur J Immunol, 2005. 35(6): p. 1670–7.

26. Dickinson, R.J. and S.M. Keyse, Diverse physiological functions for dual-specificity MAP kinase phosphatases. J Cell Sci, 2006. 119(Pt 22): p. 4607–15.

27. Fillatreau, S., Regulatory functions of B cells and regulatory plasma cells. Biomed J, 2019. 42(4): p. 233–242.

28. Gardet, A., et al., LRRK2 is involved in the IFN-gamma response and host response to pathogens. J Immunol, 2010. 185(9): p. 5577–85.

29. Sen, M., et al., COVID-19 and Eye: A Review of Ophthalmic Manifestations of COVID-19. Indian J Ophthalmol, 2021. 69(3): p. 488–509.

30. Armstrong, L., et al., In the eye of the storm: SARS-CoV-2 infection and replication at the ocular surface? Stem Cells Transl Med, 2021. 10(7): p. 976–986.

31. Fajnzylber, J., et al., SARS-CoV-2 viral load is associated with increased disease severity and mortality. Nature Communications, 2020. 11(1): p. 5493.

32. Dadras, O., et al., The relationship between COVID-19 viral load and disease severity: A systematic review. Immun Inflamm Dis, 2022. 10(3): p. e580.

33. Mathew, D., et al., Deep immune profiling of COVID-19 patients reveals distinct immunotypes with therapeutic implications. Science, 2020. 369(6508).

34. Blanco-Melo, D., et al., Imbalanced Host Response to SARS-CoV-2 Drives Development of COVID-19. Cell, 2020. 181(5): p. 1036–1045.e9.

35. Heufler, C., et al., Interleukin-12 is produced by dendritic cells and mediates T helper 1 development as well as interferon-gamma production by T helper 1 cells. Eur J Immunol, 1996. 26(3): p. 659–68.

36. Shemesh, A., et al., Differential IL-12 signaling induces human natural killer cell activating receptor-mediated ligand-specific expansion. J Exp Med, 2022. 219(8).

37. Nikkhoo, B., et al., Elevated interleukin (IL)-6 as a predictor of disease severity among Covid-19 patients: a prospective cohort study. BMC Infect Dis, 2023. 23(1): p. 311.

38. Liu, F., et al., IL-10-Producing B Cells Regulate T Helper Cell Immune Responses during 1,3-β-Glucan-Induced Lung Inflammation. Front Immunol, 2017. 8: p. 414.

39. Sun, J., et al., Transcriptomics Identify CD9 as a Marker of Murine IL-10-Competent Regulatory B Cells. Cell Rep, 2015. 13(6): p. 1110–1117.

40. Thacker, E.L., Immunology of the Porcine Respiratory Disease Complex. Veterinary Clinics of North America: Food Animal Practice, 2001. 17(3): p. 551–565.

41. Saade, G., et al., Coinfections and their molecular consequences in the porcine respiratory tract. Veterinary Research, 2020. 51(1): p. 80.

42. Davis, H.E., et al., Long COVID: major findings, mechanisms and recommendations. Nature Reviews Microbiology, 2023. 21(3): p. 133–146.

43. Ho, K.S., et al., Impact of corticosteroids in hospitalised COVID-19 patients. BMJ Open Respiratory Research, 2021. 8(1): p. e000766.

44. Närhi, F., et al., Implementation of corticosteroids in treatment of COVID-19 in the ISARIC WHO Clinical Characterisation Protocol UK: prospective, cohort study. The Lancet Digital Health, 2022. 4(4): p. e220–e234.

45. Sette, A. and S. Crotty, Adaptive immunity to SARS-CoV-2 and COVID-19. Cell, 2021. 184(4): p. 861–880.

46. Rydyznski Moderbacher, C., et al., Antigen-Specific Adaptive Immunity to SARS-CoV-2 in Acute COVID-19 and Associations with Age and Disease Severity. Cell, 2020. 183(4): p. 996–1012.e19.

47. Keep, S., et al., Identification of Amino Acids within Nonstructural Proteins 10 and 14 of the Avian Coronavirus Infectious Bronchitis Virus That Result in Attenuation In Vivo and In Ovo. J Virol, 2022. 96(6): p. e0205921.

48. Reed, L.J. and H. Muench, A SIMPLE METHOD OF ESTIMATING FIFTY PER CENT ENDPOINTS12. American Journal of Epidemiology, 1938. 27(3): p. 493–497.

49. Griffiths, J.A., et al., Detection and removal of barcode swapping in single-cell RNA-seq data. Nature communications, 2018. 9(1): p. 1–6.

50. Lun, A.T., et al., EmptyDrops: distinguishing cells from empty droplets in droplet-based single-cell RNA sequencing data. Genome biology, 2019. 20: p. 1–9.

51. L. Lun, A.T., K. Bach, and J.C. Marioni, Pooling across cells to normalize single-cell RNA sequencing data with many zero counts. Genome biology, 2016. 17: p. 1–14.

52. Haghverdi, L., et al., Batch effects in single-cell RNA-sequencing data are corrected by matching mutual nearest neighbors. Nature biotechnology, 2018. 36(5): p. 421–427.

53. Lund, S.P., et al., Detecting differential expression in RNA-sequence data using quasi-likelihood with shrunken dispersion estimates. Statistical applications in genetics and molecular biology, 2012. 11(5).

54. Lun, A.T., A.C. Richard, and J.C. Marioni, Testing for differential abundance in mass cytometry data. Nature methods, 2017. 14(7): p. 707–709.

